# Midkine and Ptprz1b act upstream of Wnt planar cell polarity to establish a midline in the developing zebrafish hindbrain

**DOI:** 10.1101/2023.11.14.566991

**Authors:** Yao Le, Kavitha Rajasekhar, Tricia Y.J. Loo, Timothy E. Saunders, Thorsten Wohland, Christoph Winkler

**Affiliations:** Department of Biological Sciences and Centre for Bioimaging Sciences, National University of Singapore, Singapore 117543, Singapore; Mechanobiology Institute, National University of Singapore, Singapore 117411, Singapore; Division of Biomedical Sciences, Warwick Medical School, University of Warwick, Coventry, CV4 7AL, United Kingdom; Department of Chemistry, National University of Singapore, Singapore 117543, Singapore

**Keywords:** Planar cell polarity, PCP, noncanonical Wnt signaling, C-division, proximity-ligation assay, fluorescence cross correlation spectroscopy, zebrafish

## Abstract

A midline in the developing central nervous system (CNS) is essential for the symmetric distribution of neural progenitors that later establish functional, bilaterally symmetric neural circuits. In the zebrafish hindbrain, a midline forms early during neurulation and requires a coordinated interplay of cell convergence and midline-crossing cell divisions (C-divisions). These two processes are controlled by the Wnt/planar cell polarity (PCP) pathway. However, upstream cues that control the timely production of PCP components remain unknown. Midkine (Mdk) and pleiotrophin (Ptn) are structurally related heparin-binding growth factors that are dynamically expressed in the developing zebrafish hindbrain. We used proximity ligation assays (PLAs) and fluorescence cross correlation spectroscopy (FCCS) *in vivo* to show that two zebrafish Mdks, Mdka and Mdkb, as well as Ptn interact with protein tyrosine phosphatase receptors type Z1, Ptprz1a and Ptprz1b, with distinct affinities. Ligand binding triggered Ptprz1b internalization and thereby determined the availability of signaling receptor on cell membranes. In zebrafish *mdka, ptn* and *ptprz1b* mutants, cell migration and convergence were significantly impaired during hindbrain neurulation. Impaired convergence led to misplaced C-divisions, defective cell polarity and consequently duplicated midlines. These duplications were rescued by overexpression of *Drosophila* Prickle, a key component of the Wnt/PCP pathway. Here, we provide evidence that zygotic Mdka controls the distribution of maternally provided Ptprz1b, which in turn is needed for transcription of zebrafish *prickle1b*. Our findings thus reveal a role for Mdka and Ptprz1b upstream of Wnt/PCP to coordinate neural plate convergence, neural progenitor positioning and midline formation.

## INTRODUCTION

During vertebrate neurulation, a single midline structure is established in the central nervous system (CNS) to allow bilateral symmetric distribution of neurons as well as formation of a ventricle or lumen (Clarke, 2009). Establishment of this midline is achieved through a complex interplay of convergent extension (CE) and midline-crossing cell divisions (C-divisions) (Clarke, 2009). In zebrafish, CE narrows the neuroepithelium into a solid neural keel structure, to position neural progenitor cells close to the presumptive midline where they undergo C-divisions (Concha & Adams, 1998; Hong & Brewster, 2006; Tawk *et al*, 2007). After C-division, one daughter cell remains in the original half of the neural keel, while the other crosses the presumptive midline to intercalate into the contralateral side. This leads to a bilateral distribution of two descendants of the same neural progenitor (Ciruna *et al*, 2006; Tawk *et al*., 2007). C-divisions depend on a correct establishment of anteroposterior and apicobasal polarity (Buckley *et al*, 2013; Ciruna *et al*., 2006; Tawk *et al*., 2007). While anteroposterior polarity is induced by noncanonical Wnt/planar cell polarity (PCP) signaling that drives CE and allows contralateral intercalation of daughter cells after C-divisions (Ciruna *et al*., 2006; Tawk *et al*., 2007), apicobasal cell polarity ensures that C-divisions occur at correct medial positions along the mediolateral axis (Tawk et al., 2007; Buckley et al., 2013). Cues that act upstream of Wnt/PCP to control cell polarity and positioning of C-divisions, however, remain poorly understood (Hirano *et al*, 2022).

Midkine (MDK) and pleiotrophin (PTN) are highly conserved heparin-binding growth factors implicated in various processes in the nervous system, including neuronal maturation (Tang et al., 2019), neurite outgrowth (Kaneda et al., 1996b; Kinnunen et al., 1996), cell migration (Maeda and Noda, 1998), and metastasis (Olmeda et al., 2017; Qin et al., 2017; Shi et al., 2017). Earlier studies proposed chondroitin sulfate side chains of the protein tyrosine phosphatase receptor type Z1 (PTPRZ1) as possible binding sites for human MDK and PTN (Maeda *et al*, 1999; Maeda *et al*, 1996), but evidence supporting such interactions *in vivo* is lacking. Cell culture and cancer studies suggested that downstream targets of MDK/PTN-PTPRZ1 signaling play a role in cytoskeletal remodeling and cell adhesion (Kuboyama *et al*, 2015; Meng *et al*, 2000; Qin *et al*, 2017; Shi *et al*, 2017). In glioma, PTN-PTPRZ1 signaling facilitates tumor invasion by activating Rho/ROCK signaling and triggering neural stem cell-like mitotic behavior (Bhaduri *et al*, 2020; Qin *et al*., 2017). Recent studies also revealed that PTN-PTPRZ1 regulates mitosis of human outer radial glial (oRG) cells, to maintain their stemness (Bhaduri *et al*., 2020; Pollen *et al*, 2015). In mouse, *Mdk, Ptn*, and *Ptprz1* are dynamically expressed in the CNS during neurulation (Fan *et al*, 2000; Shintani *et al*, 1998), but their knockout only induced subtle defects (Amet *et al*, 2001; Harroch *et al*, 2000; Nakamura *et al*, 1998). A double knockout of *Mdk* and *Ptn*, on the other hand, led to increased lethality before E14.5 (Muramatsu *et al*, 2006) suggesting redundant but indispensable functions for MDK and PTN. Unlike mammalian genomes that contain single *MDK, PTN* and *PTPRZ1* genes, a teleost-specific genome duplication resulted in two *mdk* (*mdka* and *mdkb*), one *ptn*, and two *ptprz1* (*ptprz1a* and *ptprz1b*) genes in zebrafish (van Eekelen *et al*, 2010; Winkler *et al*, 2003; Winkler & Yao, 2014). These additional paralogs are targets for possible neo- or sub-functionalization, increasing the complexity of ligand-receptor interactions that require precise spatiotemporal control. Zebrafish *mdka/b, ptn* and *ptprz1a/b* are dynamically expressed in the developing CNS, but their exact functions remain to be identified (Chang *et al*, 2004; Schafer *et al*, 2005; van Eekelen *et al*., 2010; Winkler & Moon, 2001b).

In the present study, we show that Mdka, Mdkb and Ptn bind to Ptprz1a and Ptprz1b *in vivo* with different affinities. Our findings suggest that Mdka and Ptn diffuse over long distances in embryos to interact with Ptprz1b. Confocal time-lapse imaging of hindbrain rhombomeres revealed distinct transient midline defects in *mdka, ptn* and *ptprz1b* mutants.

These were caused by defective CE, which placed neural progenitor cells at aberrant positions to perform C-divisions resulting in duplicated or ectopic midlines. These midline defects were partially rescued by overexpression of *Drosophila* Prickle, a Wnt/PCP core component. As *prickle1b* was down-regulated in *mdka* and *ptprz1b* mutants, this suggests that Mdka/Ptn-Ptprz1b acts upstream of noncanonical Wnt/PCP signaling to control midline formation in zebrafish rhombomeres.

## RESULTS

### Mdka, Mdkb and Ptn interact with Ptprz1 with different affinities *in vivo*

Zebrafish *mdka, mdkb* and *ptn* are dynamically expressed during hindbrain morphogenesis (**Fig 1A-C**). Fluorescent *in situ* hybridization (FISH) at 14 hours post fertilization (hpf) revealed distinct *mdka* and *mdkb* expression in hindbrain rhombomeres 1 (r1) to r6, with decreasing expression from anterior to posterior (**Fig 1A, C**). *ptn* transcription was restricted to r5 and r6 and overall levels appeared to be lower when compared to *mdka* and *mdkb* (**Fig 1A, C**). Thus, the three structurally related ligands (70.83% amino acid identity for Mdka and Mdkb, 50.00% for Mdka/Ptn and 47.83% for Mdkb/Ptn, BLAST) are expressed in distinct, largely overlapping domains in rhombomeres during neurulation.

**Figure 1.**
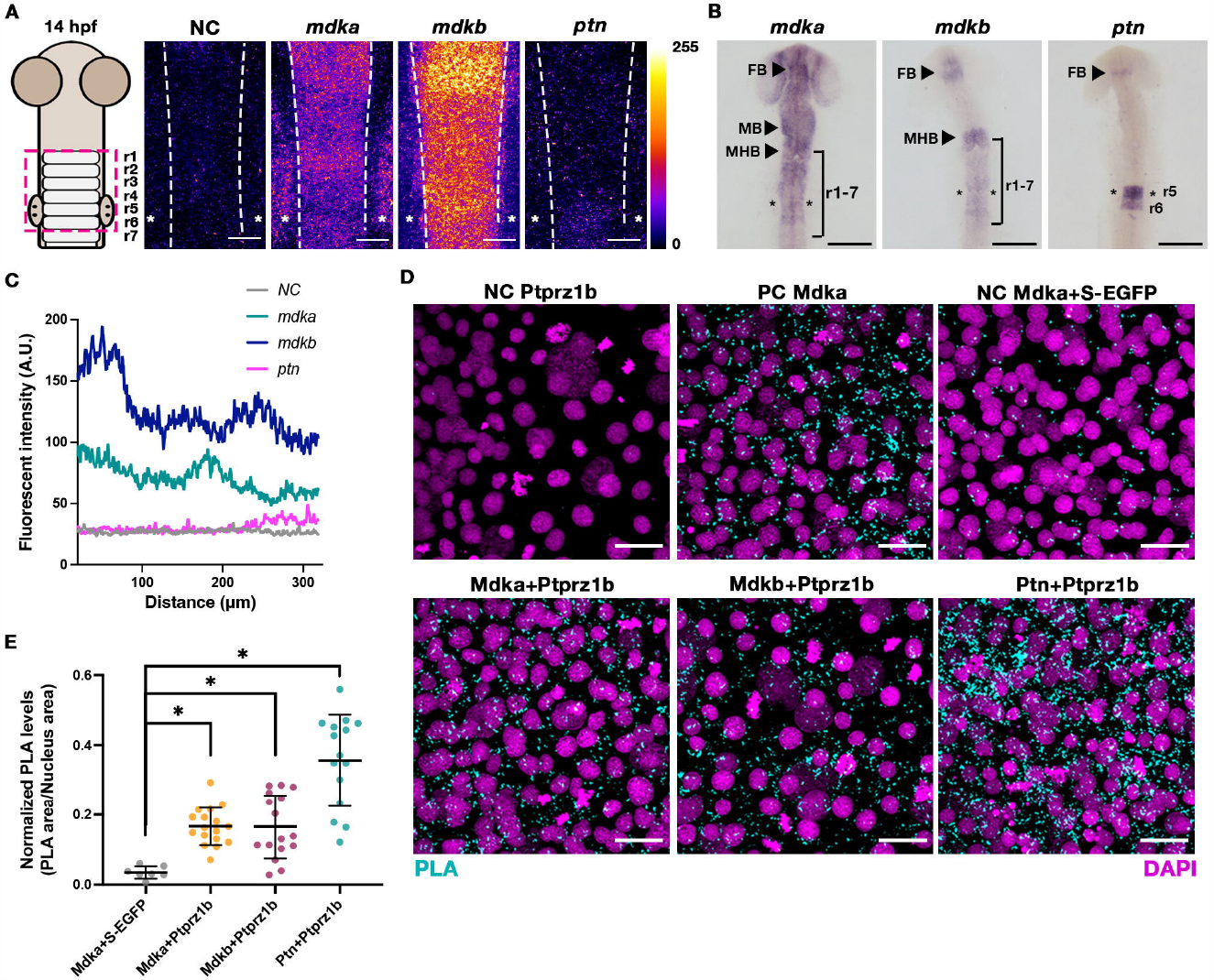
Expression and receptor interaction of zebrafish Mdka, Mdkb and Ptn. (**A**) Fluorescent RNA *in situ* hybridization (FISH) of *mdka, mdkb* and *ptn* expression in wildtype embryos at 14 hpf. Left: Schematic diagram of hindbrain organization, dorsal view with dashed box indicating rhombomere region analyzed by FISH. NC = negative control (*mdka* sense probe). Representative maximum-intensity-projection (MIP) images showing dorsal views and pseudo-coloured in fire LUT with an intensity range from 0 to 255 by ImageJ. Dotted lines delineate lateral edges of rhombomeres, white asterisks indicate position of otic vesicles at r5. Scale bars = 50 μm. (**B**) Whole mount RNA *in situ* hybridization of *mdka, mdkb* and *ptn* expression at 18 hpf. Dorsal views of head regions with rhombomere region r1-7. FB, forebrain; MHB, midbrain-hindbrain boundary; r1-7, rhombomeres 1 to 7. Scale bars = 200 μm. (**C**) Mean fluorescent intensity of FISH signals in (**A**) along anteroposterior axis of rhombomeres. (**D**) Representative MIP images after proximity ligation assay (PLA) of controls and different pairs of ligands (Mdka, Mdkb, Ptn) and receptor (Ptprz1b). Embryos injected with *HA-ptprz1b* mRNA alone served as the negative control (NC). Positive controls (PC) are *mdka*-*MYC* mRNA-injected embryos using two pairing secondary antibodies that recognize the same primary antibody. Embryos co-injected with *mdka-MYC* and *secreted EGFP* (*secEGFP*) mRNA served as random collision controls. PLA signals are represented in cyan, and DAPI in magenta. Scale bars = 20 μm. (**E**) Statistical analysis of PLA levels normalized to DAPI signals after thresholding (PLA/DAPI area ratios). Data are presented as scattered dots with mean ± SD. Statistical analysis was performed on Estimation Stats (https://www.estimationstats.com) to compare each dataset with the random collision control. *P* values are calculated from a two-sided permutation *t*-test under a CI of 95%. Asterisks indicate statistical significance (*P* < 0.05).

Previous *in vitro* findings proposed PTPRZ1 as candidate receptor for MDK and PTN (Maeda *et al*., 1999; Maeda *et al*., 1996), but their binding dynamics *in vivo* remained unclear. To test whether zebrafish Mdka/b and Ptn bind to Ptprz1a/b *in vivo*, we used proximity ligation assays (PLAs), visualized interaction events and compared relative binding affinities of various ligand-receptor combinations (**Fig S1A**). PLA detects targets that are in close physical proximity of less than 30 nm (Soderberg *et al*, 2006). Injection of mRNAs encoding MYC-tagged Mdka/Mdkb/Ptn ligands and HA-tagged Ptprz1a/Ptprz1b receptors resulted in homogenously distributed expression in early embryos, similar to the situation of maternally provided endogenous proteins. mRNAs were injected into wildtype (WT) embryos at 1-cell stage and PLA was performed at 80% epiboly (**Fig S1B**). This identified interaction of Mdka, Mdkb and Ptn with Ptprz1b with different affinities (**Fig 1D-E**). Signal levels were significantly higher than those from random collision of Mdka and secEGFP (**Fig 1D-E**) (*P* < 0.05, CI = 95%). Normalized PLA signal ratios (PLA area/nucleus area) were highest for Ptn-Ptprz1b (mean ± SD = 0.36 ± 0.13), followed by Mdkb-Ptprz1b (mean ± SD = 0.17 ± 0.09) and Mdka-Ptprz1b (mean ± SD = 0.17 ± 0.05) (**Fig 1D-E**). PLA with the co-ortholog Ptprz1a also revealed significant binding (**Fig S1C-E**). Mdka showed comparable binding towards Ptprz1a and Ptprz1b, while Mdkb and Ptn displayed higher binding affinities towards Ptprz1b than to Ptprz1a (**Fig S1F**). This indicates that Mdka, Mdkb and Ptn bind to Ptprz1 receptors *in vivo* and suggest distinct binding preferences when different ligands and receptors are expressed at similar levels. For confirmation, we utilized dual color fluorescence cross-correlation spectroscopy (DC-FCCS) for real-time measurements of Ptn and Ptprz1b interactions. There was a significantly increased relative cross-correlation value between Ptn and Ptprz1b (Q = 0.70 ± 0.19, mean ± SD) when compared to a negative control (Q_neg_ = 0.14 ± 0.06, mean ± SD) and a comparable value to a positive control (Q_pos_ = 0.67 ± 0.08, mean ± SD) (**Fig S1G-J**). This confirmed that Ptn binds to Ptprz1b binding in real time *in vivo*.

### *mdka, mdkb, ptn* and *ptprz1b* mutants exhibit distinct midline defects in forming rhombomeres

To test the roles of *mdka, mdkb, ptn* and *ptprz1b* in hindbrain morphogenesis, CRISPR-Cas9 mutants were generated and analyzed (**Fig S2**). In *PMT-mEGFP* mRNA injected WT embryos (N = 5), confocal time-lapse imaging of the neural rod revealed establishment of a single midline between 16 to 17 hpf, as previously reported (Tawk *et al*., 2007) (**Fig 2A, A’, E, E’;Movie S1**). In contrast, stage-matched MZ *ptprz1b* mutant embryos (N = 6) formed two, ectopically placed midlines starting from 16 hpf (**Fig 2C-D, C’-D’, F, F’; Fig S3A-B, E, A’-B’, E’;Movies S2, S3**), which was most obvious at the level of r2 and r3 (**Fig 2K-L**). Eventually, the two midlines merged at 17 to 18 hpf (**Fig 2D, D’**), except in one case where duplicated midlines persisted throughout the imaging period (**Fig S3A-B, A’-B’**). In orthogonal views, ectopic midlines were most pronounced in the dorsal neural rod (**Fig 2E-F, E’-F’; Fig S3E, E’**). MZ *mdka* mutants (N = 6) also had midline defects with high penetrance in r1, r2, r4 and r5, but not r3 (**Fig 2G-H, G’-H’, K; Fig S3C, C’;Movies S4, S5**). These defects appeared to be more severe than in MZ *ptprz1b* mutants, as only 3 out of 6 analyzed MZ *mdka* embryos displayed a recovery at 18 hpf (**Fig 2G-H, G’-H’; Table 1**), while in other embryos, ectopic midlines persisted beyond 22 hpf (**Fig S3C, C’**). Together, the similar rhombomere phenotypes in *mdka* and *ptprz1b* mutants suggested that Mdka acts through Ptprz1b to control formation of the hindbrain midline. MZ *ptn* mutants (N = 6), on the other hand, exhibited less severe midline defects with lower penetrance (**Fig 2I-J, I’-J’; Fig S3D, D’; Table 1**). Ectopic midlines were obvious at r5, r6 or r7, and merged at 18 hpf (N = 2/3) (**Fig 2I-J, I’-J’; Fig S3D, D’**). In r5 and r6, the penetrance of phenotypes was similar in MZ *ptn* mutants and MZ *ptprz1b* mutants (**Fig 2L**), suggesting a functional Ptn-Ptprz1b interaction predominantly in posterior rhombomeres. Next, junctional F-actin distribution was assessed in MZ *ptprz1b* and MZ *mdka* mutants. The hindbrain midline structure is established by apical surfaces of two bilaterally positioned, opposing neural progenitors (Geldmacher-Voss et al., 2003). In WT embryos, F-actin staining was enriched along these surfaces indicating intact apical cortex organization at the midline (**Fig 2M, M’, P**). In MZ *ptprz1b* mutants, in contrast, 13 out of 17 embryos showed aberrant F-actin accumulation with two evident peaks of phalloidin staining in orthogonal views (**Fig 2N, N’, Q; Fig S3F, H, F’**). Nine out of these 13 embryos showed ectopic midlines most predominantly in r2 and r3 (**Fig 2N, N’, Q**), while the remaining four exhibited a severe phenotype with complete failure of convergence (**Fig S3F, F’, H**). This confirmed the findings of time-lapse confocal imaging (**Table 1**). For MZ *mdka* mutants, 25 out of 31 embryos showed two peaks of phalloidin signal indicative for midline duplications (**Fig 2O, O’, R**). Five of these 25 mutant embryos failed to converge and had severe midline defects (**Fig S3G, G’, I**). This suggests that ectopic midline formation in MZ *ptprz1b* and MZ *mdka* mutants was accompanied by ectopic F-actin accumulation as possible indicator of impaired cell polarity. MZ *mdkb* mutants, in contrast, formed a single midline, however with a delay (**Fig S4A-F**) consistent with the idea of a possible functional divergence of Mdka/Ptn and Mdkb. The nuclei were also aberrantly positioned in MZ *ptprz1b* and MZ *mdka* mutant rhombomeres. In WT, as indicated by DAPI stained nuclei, cells were regularly stacked along the anteroposterior and dorsoventral axes in each half of the neural rod. Most nuclei were excluded from the midline, while those found within the midline were mostly mitotic (**Fig 2M, M’, P**). In contrast, nuclei in MZ *ptprz1b* and MZ *mdka* mutants were mostly positioned in a random manner along the mediolateral axis (**Fig 2N-O, N’-O’, Q-R**). Orthogonal views showed more than one cell along the mediolateral axis and ectopic cell accumulation in the middle of the dorsal neural rod (**Fig S3L-Q**). In DAPI histograms, two signal peaks were evident along the mediolateral axis of WT hindbrains with nearly no nuclei in the middle (**Fig S3L, O**). In contrast, MZ *ptprz1b* and MZ *mdka* mutants showed multiple signal peaks (**Fig S3M-N, P-Q**). Together, both live imaging and histochemical analysis of MZ mutants indicated aberrant positioning of neural progenitor nuclei and apical surfaces suggesting that Mdka/Ptprz1b signaling is crucial for correct positioning of cells in order to establish a midline in rhombomeres.

**Table 1.** Quantification of midline phenotypes observed by time-lapse imaging.

| Genotype<br>Phenotype | WT | MZ *ptprz1b* | MZ *mdka* | MZ *ptn* |
| --- | --- | --- | --- | --- |
| Single midline | 5 | - | - | 3 |
| Ectopic midlines (mild) | - | 5 | 3 | 2 |
| Ectopic midlines (severe) | - | 1 | 3 | 1 |
| Rescued at 18 hpf | - | 5 | 3 | 2 |
| Persistent after 18 hpf | - | 1 (r1-7) | 3 (r2-3) | 1 (r1-7) |
| Total | 5 | 6 | 6 | 6 |

**Figure 2.**
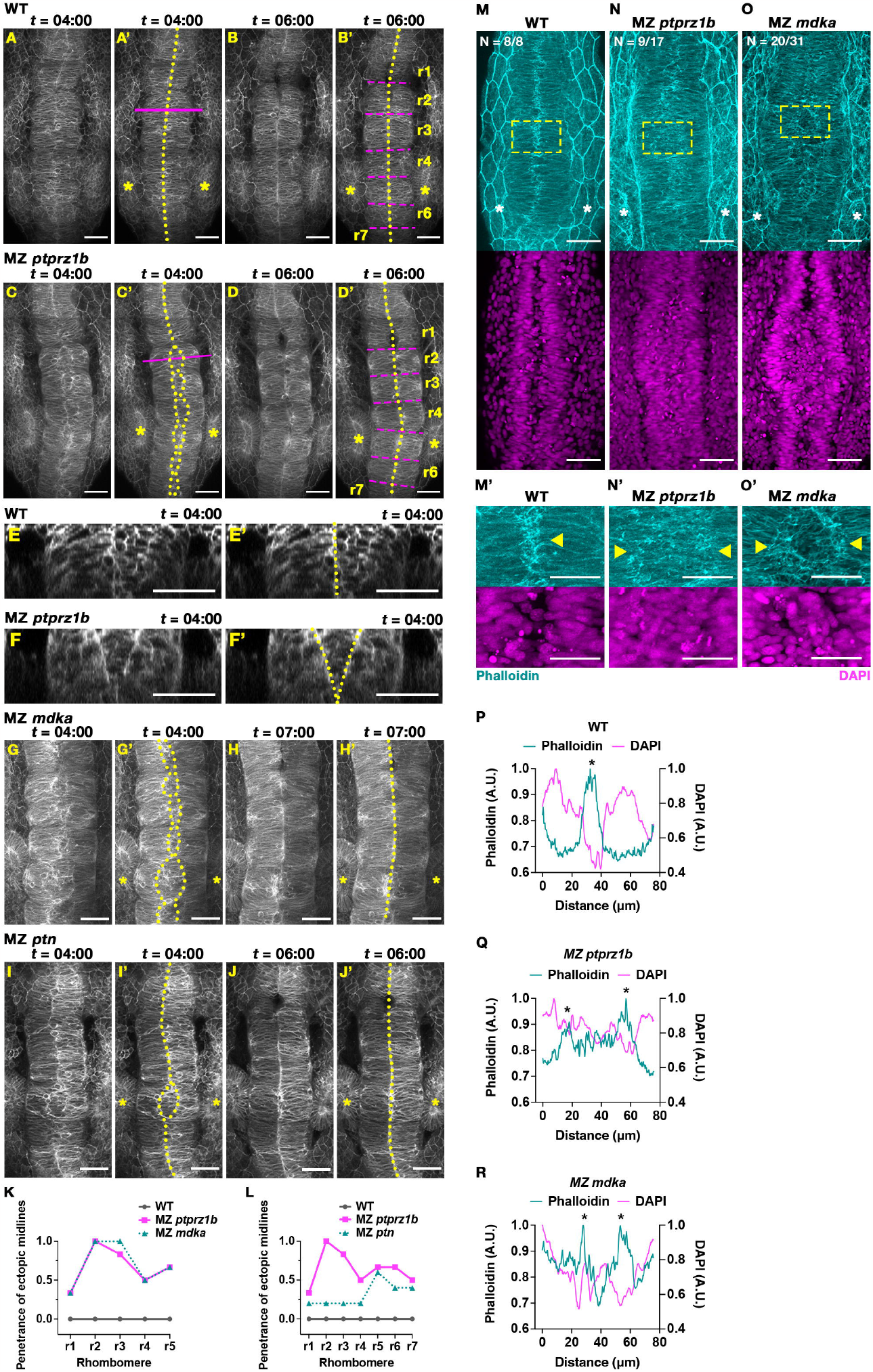
MZ *ptprz1b*, MZ *mdka* and MZ *ptn* mutants exhibit transient ectopic midline formation in rhombomeres. (**A-B**) Representative MIP still images in dorsal view taken from confocal time-lapse analysis at 4 and 6 h of time-lapse showing normal midline formation in rhombomeres of WT embryos. Imaging was done from approx. 13 hpf with 4 min intervals. PMT-mEGFP was injected to mark cell membranes. images. Elapsed time (*t*) is displayed as hours:minutes (hh:mm). Midline structures are delineated by dotted lines, magenta solid line indicates position of views shown in (**E, E’**). Magenta dotted lines delineate rhombomere boundaries. Asterisks label position of otic vesicles at r5. Scale bar = 50 μm. (**C-D**) Transient ectopic midline in MZ *ptprz1b* mutants. Magenta solid line indicates position of views shown in (**F, F’**). Scale bar = 50 μm. (**E-F**) Computational orthogonal view of r2 in WT and MZ *ptprz1b* mutant. Yellow dotted lines delineate midline structures. Scale bar = 50 μm. (**G-H**) Transient ectopic midline in MZ *mdka* mutant. Still images taken from confocal time-lapse analysis at 4 and 7 h. Scale bar = 50 μm. (**I-J**) Transient ectopic midline in MZ *ptn* mutant. Scale bar = 50 μm. (**K-L**) Quantification of penetrance of ectopic midline phenotype in each rhombomere in WT (N = 5), MZ *ptprz1b* (N = 6), MZ *mdka* (N = 6) and MZ *ptn* mutants (N = 6). The penetrance of ectopic midlines is calculated by dividing the number of embryos with ectopic midlines at a given rhombomere by the total number of embryos. (**M-O**) Representative MIP images of Phalloidin stained F-actin (cyan) and DAPI stained nuclei (magenta) in rhombomere region of WT, MZ *ptprz1b* and MZ *mdka* mutant embryos at 17 hpf. Boxes indicate region shown in (**M’-O’**). Scale bar = 50 μm (**M-O**) and 30 μm (**M’-O’**). Arrowheads indicate F-actin accumulation. (**P-R**) Respective mean fluorescent intensity histogram of Phalloidin (cyan) and DAPI (magenta) along mediolateral axis of (**M’-O’**). Asterisks highlight aggregation of Phalloidin-stained F-actin.

### Mdka and Ptn diffuse over long distances to interact with Ptprz1b

We noted that midline defects in MZ *mdka* mutants were transient, and in most cases had fully recovered at 17 hpf. This opened the possibility that Ptn had compensated for the loss of Mdka activity. However, *ptn* expression is restricted to r5 and r6 and thus requires long-ranged transport to reach r1 to r4. Previous studies had reported that MDK and PTN act at considerable distances from their source (Olmeda *et al*, 2017; Qin *et al*., 2017; Shi *et al*., 2017) implying long range diffusion despite efficient heparin-binding. Importantly, such diffusion dynamics of MDK/PTN *in vivo* remained unknown.

To visualize long range Mdk and Ptn diffusion *in vivo*, we injected and expressed epitope-tagged Mdka-MYC/Ptn-MYC in source cells away from mEGFP-Ptprz1b expressing cells in early zebrafish embryos (**Fig 3A**). Immunostaining revealed that Mdka and Ptn were present across the entire blastula at 4.3 hpf (**Fig 3B-E**). Unexpectedly, fluorescence signals for Mdka and Ptn were considerably lower inside the mEGFP-Ptprz1b expression domain suggesting that Ptprz1b either hindered diffusion or facilitated a decay of the ligands (**Fig 3D-E**). PLA also demonstrated substantial Ptn binding to Ptprz1b at a distance (**Fig 3F-G**). This opened the possibility that Mdka and Ptn act redundantly even though expressed in distant rhombomeres.

**Figure 3.**
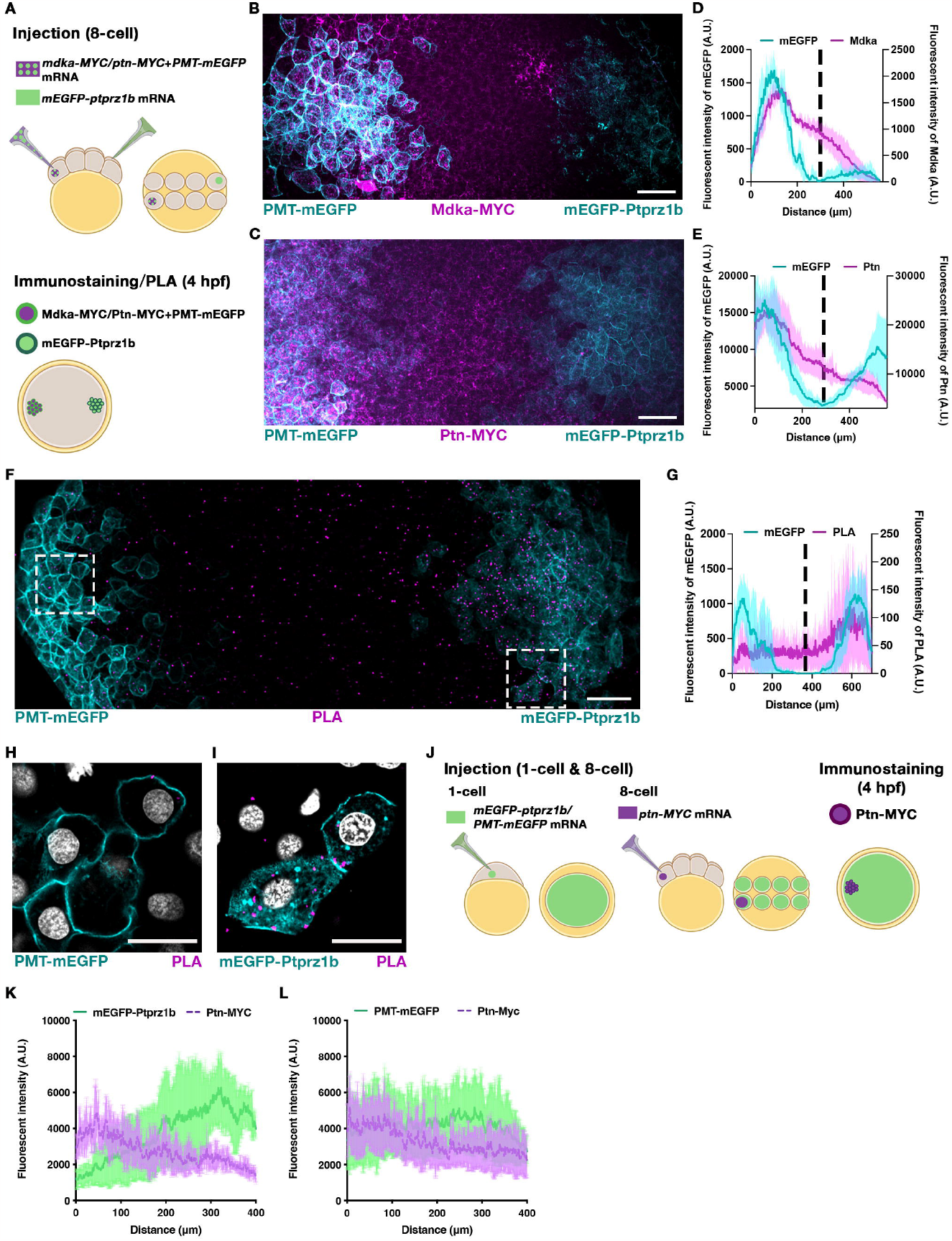
Diffusion of Mdka and Ptn triggers Ptprz1b receptor internalization. (**A**) Experimental outline to visualize Mdka and Ptn diffusion. At 8-cell stage, two distant cells were injected, one with a mix of either *mdka-MYC* or *ptn-MYC* with *PMT-mEGFP* mRNAs, the other with *mEGFP-ptprz1b* mRNA. At 4 hpf, two distinct mEGFP expressing domains could be identified. One was the PMT-mEGFP domain with mEGFP present exclusively on cell membranes, the second was the mEGFP-Ptprz1b domain with mEGFP on membranes and in cytoplasm. Mdka or Ptn were secreted from the PMT-mEGFP domain and diffused across the blastula to reach the mEGFP-Ptprz1b domain. (**B-C**) Representative animal-pole MIP images of blastula embryos after diffusion of Mdka (**B**, magenta) or Ptn (**C**, magenta), secreted from PMT-mEGFP expressing cells (cyan) on the left, towards mEGFP-Ptprz1b expressing cells (cyan) on the right. Scale bar = 50 μm. (**D-E**) Respective mean fluorescent intensity profiles of (**B-C**) shown as mean ± SD. Dashed lines separate domains with PMT-mEGFP expressing cells (left) from those with mEGFP-Ptprz1b expressing cells (right). (**F**) Representative MIP images of PLA analysis obtained by setup shown in (**A**). PLA signals are coloured in magenta, mEGFP signals in cyan. Scale bar = 50 μm. (**G**) Respective mean fluorescent intensity profile of(**F**) in the form of mean ± SD. (**H-I**) High magnification, single plane images of selected regions boxed in (**F**). DAPI stained nuclei are shown in gray. Scale bar = 20 μm. (**J**) Outline of 1- and 8-cell injections to study Ptn diffusion in Ptprz1b expressing tissue. Embryos at 1-cell stage were injected with *mEGFP-ptprz1b* or *PMT-mEGFP* mRNA to create homogenous distribution of mEGFP-Ptprz1b or PMT-mEGFP at 4 hpf. mRNA encoding Ptn-MYC was injected into a single cell at 8-cell stage to generate a clone of source cells. At 4 hpf, Ptn was secreted from source cells and diffused through tissues with homogenous mEGFP-Ptprz1b or PMT-mEGFP expression. Source cells were identified by bright cytoplasmic Ptn-MYC puncta. (**K-L**) Mean fluorescent intensity profile of mEGFP-Ptprz1b or PMT-mEGFP and Ptn-MYC in *mEGFP-ptprz1b* injected embryos (**K**) (N =4), or PMT-mEGFP injected embryos (**L**) (N = 5) along the left-right axis. Datasets are presented as mean ± SD. Illustrations were created with BioRender.com
.

Importantly, we noticed abundant PLA signals inside cells close to the plasma membrane suggesting a possible internalization of ligand-bound Ptprz1b receptor complexes (**Fig 3H-I**). To confirm this, we expressed Ptn-MYC in confined cell clones of zebrafish blastulae that ubiquitously expressed mEGFP-Ptprz1b (**Fig 3J**). We found mutually exclusive distribution of Ptn-MYC and mEGFP-Ptprz1b in line with the idea that locally restricted ligand secretion caused efficient receptor internalization (**Fig 3K; Fig S5A**). Importantly, the reduction of mEGFP-Ptprz1b near the ligand source was not triggered by concurrent internalization of Ptn-MYC with other membrane proteins as Ptn-MYC did not reshape the homogenous distribution of a PMT-mEGFP control (**Fig 3L; Fig S5B**).

Our results are consistent with long-ranged diffusion of Mdk and Ptn, with their range modulated by receptor binding. We tested whether this conclusion is consistent with a reaction-diffusion model of gradient formation, as has been shown for other long-ranged morphogens (Kruse *et al*, 2004; Kuhn *et al*, 2022; Wang *et al*, 2016). We incorporated diffusion of Ptn with local receptor binding, with the spatial range defined by the profiles in Fig 3D, E, G (Methods). We found that a simple model with five parameters was able to reproduce our experimental profiles (**Fig S6**).

Together, these findings suggest that transport of Mdk and Ptn is consistent with effective diffusion over long distances in embryos with profile extent modulated by binding to receptors on distant rhombomeres. Such binding triggers internalization of ligand-receptor complexes, which consequently alters receptor availability and thus defines the temporospatial pattern of ligand diffusion and receptor signaling.

As *ptprz1b* showed high maternal but only scarce zygotic expression (**Fig S7E-F**), we next tested whether ectopic midlines are caused by a loss of maternal Ptprz1b contribution. MZ *ptprz1b* mutant females were crossed with WT males, and 9 out of 11 progenies recapitulated the ectopic midline phenotype as observed in MZ *ptprz1b* mutants (**Fig S7A, B**). In contrast, embryos from reverse crosses displayed normal midlines, indicating that ectopic midlines were caused by depletion of maternal *ptprz1b* rather than insufficient zygotic *ptprz1b* (**Fig S7A, C**). In comparison, embryos lacking only maternal *mdka* (M *mdka*) exhibited normal midline morphology (**Fig S7A, D**). Together, this suggests that a combination of maternal Ptprz1b and zygotic Mdka controls midline formation in rhombomeres.

### Misplaced and disorientated cell divisions in MZ *ptprz1b* and MZ *mdka* mutant rhombomeres

In zebrafish rhombomeres, the establishment of anteroposterior cell polarity facilitates midline formation by coordinating convergent extension (CE) cell movements and midline-crossing cell divisions (C-divisions) (Ciruna *et al*., 2006). In MZ *ptprz1b* and MZ *mdka* mutants, the maximal mediolateral widths of r2, r3 and r4 were significantly increased at 15 hpf, suggesting deficient CE (**Fig 4A-C**). In contrast, MZ *mdkb* mutants revealed a decreased width of r2 at 15 hpf (**Fig S4G-I**). Earlier studies had shown that impaired CE leads to misplacement of neural progenitor cells at more lateral positions resulting in ectopic C-divisions (Tawk *et al*., 2007). Confocal time-lapse imaging of MZ *ptprz1b* and MZ *mdka* mutants revealed that most mitotic events likewise occurred more laterally in mutants (**Fig 5A-B**). This suggested that ectopic midlines in mutants resulted from defective neural keel convergence and consequently misplaced C-divisions. Medially positioned C-divisions facilitate the intercalation of one of the daughter cells into the contralateral half of the neural keel (Tawk *et al*., 2007). Misplaced C-divisions, in contrast, cause a failure in contralateral intercalation, leading to ectopic cell accumulation and midline duplications (Ciruna *et al*., 2006; Tawk *et al*., 2007). Through single cell tracking, WT cells were found to display characteristic C-divisions that deposited two daughter cells in a mirror-symmetrical fashion into each half of the neural keel (**Fig 5D;Movie S6**). On the other hand, neural progenitor cells in MZ *ptprz1b* (**Fig 5E;Movie S7**) and MZ *mdka* mutants (**Fig 5F;Movie S8**) exhibited impaired contralateral intercalation. While daughter cells close to the lateral boundary integrated properly proximally, the other daughter cell was unable to intercalate into the contralateral half of the neural keel. These cells were principally mobile and able to cross the presumptive ectopic midline but then became stagnant and remained in the middle of the neural keel with stochastic and undirected movements (**Fig 5E-F**). Ectopic cell accumulation between the two forming midlines was thus likely due to a failure of contralateral intercalation after misplaced C-division, with two midlines emerging at sites of ectopic C-divisions (**Fig 5E-F**). Continuous cell tracking showed that the stagnant daughter cells in both mutants eventually reinitiated cell migration and intercalated into the neural rod as ectopic midlines converged at 18 hpf (**Fig 5E-F;Movies S7, S8**). These data suggest that ectopic midlines are rescued by re-intercalation of ectopic cells in both MZ *ptprz1b* mutants and MZ *mdka* mutants.

**Figure 4.**
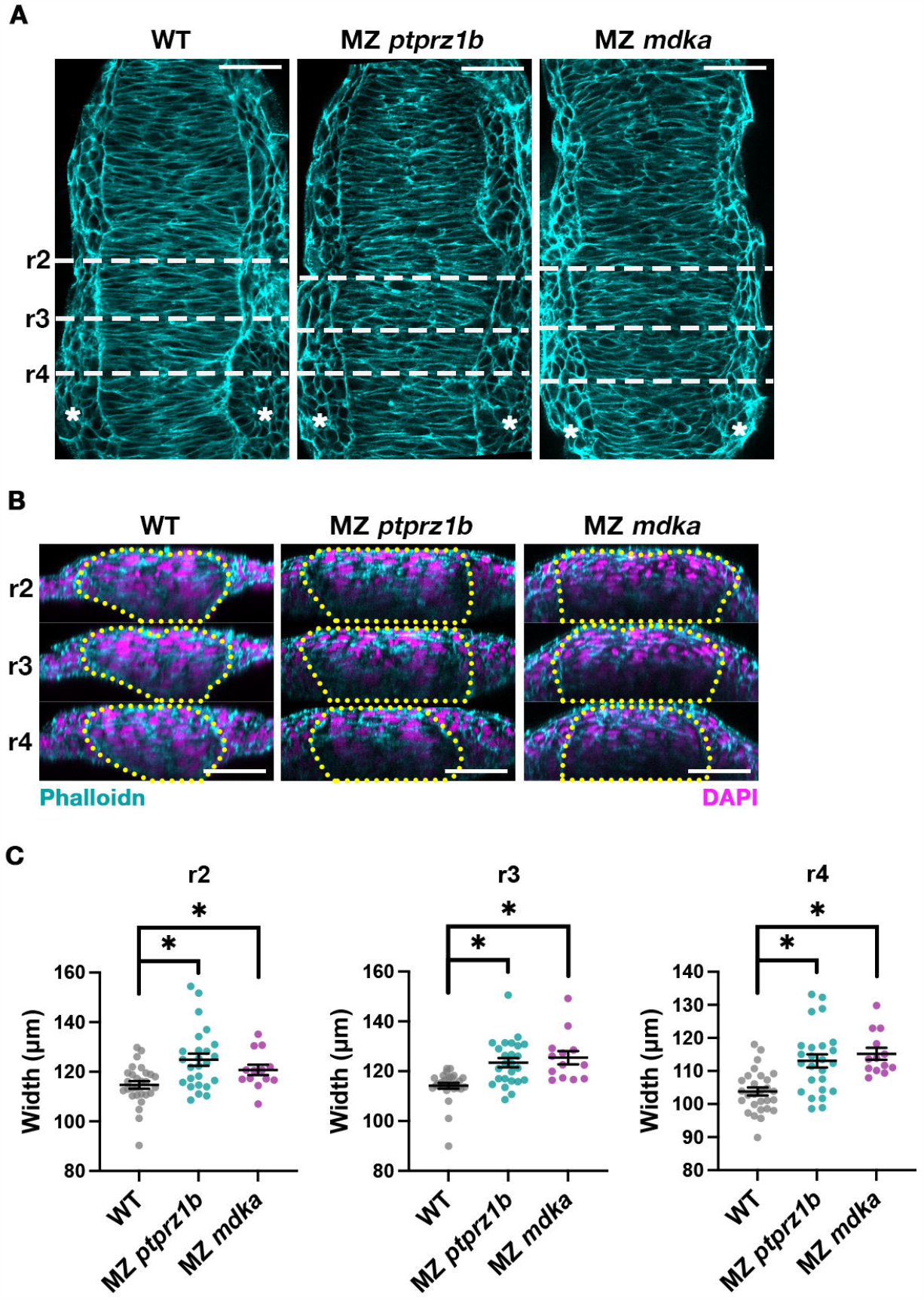
MZ *ptprz1b* and MZ *mdka* mutants exhibit delayed neural keel convergence. (**A**)Representative confocal images (single-plane, dorsal views) of Phalloidin-stained hindbrains in 15 hpf WT (N = 28), MZ *ptprz1b* (N = 25), and MZ *mdka* (N = 13) embryos. Asterisks mark positions of otic vesicles, dotted lines indicate positions of orthogonal views shown in (**B**). Scale bars = 50 μm. (**B**)Reconstructed orthogonal views of embryos shown in (**A**). Phalloidin-stained F-actin is pseudo-coloured in cyan, and DAPI-stained nuclei in magenta. Scale bars = 50 μm. (**C**)Quantification of the maximum width measured for individual r2, r3 and r4. Data are shown as scattered dots. Mean ± SD of each dataset is indicated by line and error bar. Statistical comparison was performed between WT (N = 28) and MZ *ptprz1b* (N = 25) or MZ *mdka* (N = 13) on Estimation Stats (https://www.estimationstats.com). A two-sided permutation *t*-test was performed to calculate *P* values under a CI of 95%. Asterisks indicate statistical significance (*P* < 0.05).

**Figure 5.**
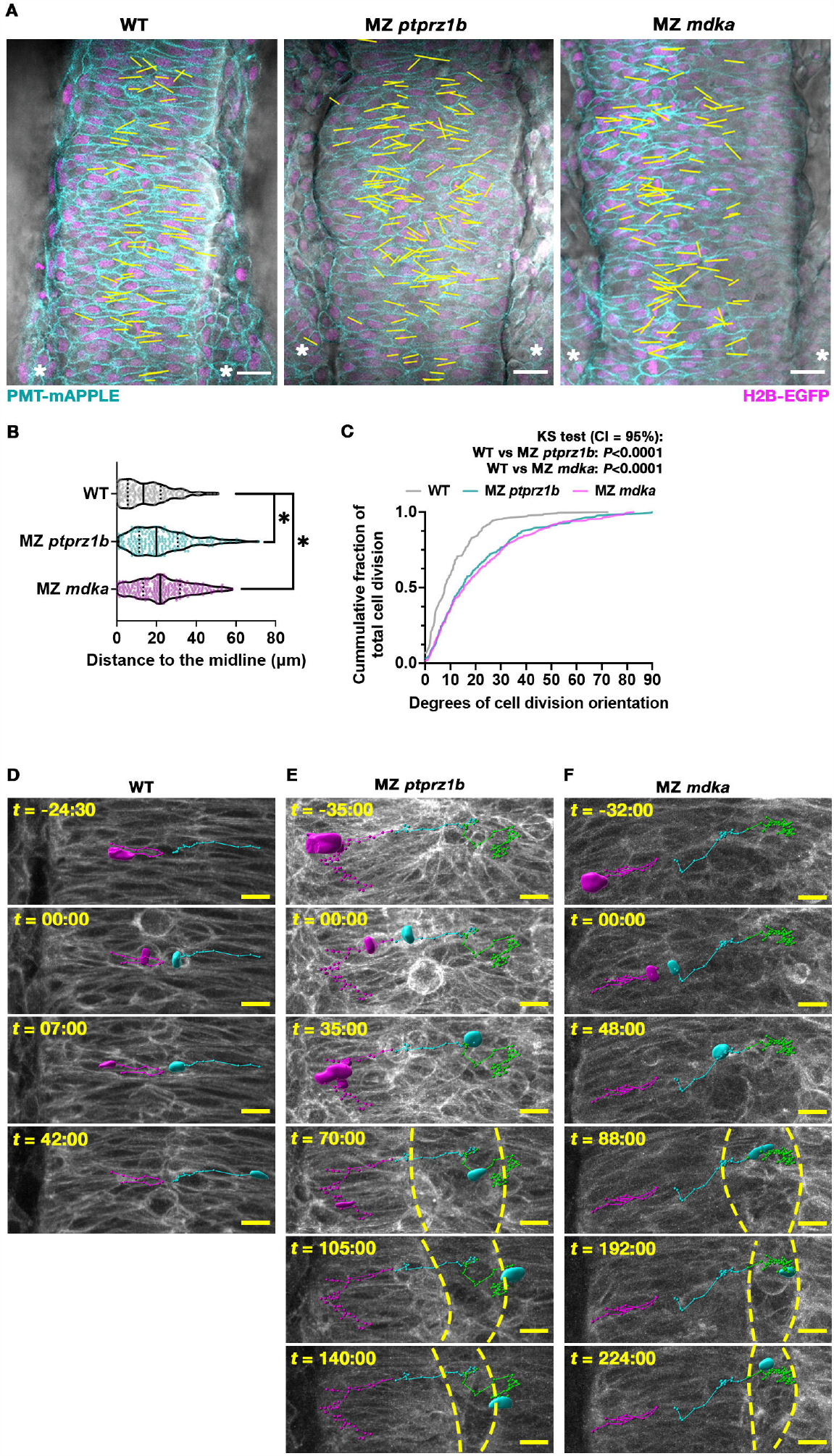
MZ *ptprz1b* and MZ *mdka* mutant cells show misplaced and disoriented C-divisions and a failure in contralateral intercalation. (**A**)Representative single-plane still images (dorsal views) taken from time-lapseMovies of WT (N = 4), MZ *ptprz1b* (N = 3) and MZ *mdka* mutants (N = 3). PMT-mApple (cyan) labels cell membranes, H2B-EGFP (magenta) marks nuclei and DIC is shown in gray. Yellow lines indicate XY-position and direction of C-divisions superimposed from Z-planes and timepoints of interests. Lines are drawn by linking the respective middle points of two separating pairs of chromosomes at telophase. Scale bar = 20 μm. (**B**)Quantification of relative distances of C-divisions to middle of neural keel in WT (N = 4), MZ *ptprz1b* (N = 3) and MZ *mdka* mutants (N = 3) mutants. A total of 215, 201 and 269 C-division events were identified and analyzed in WT, MZ *ptprz1b* and MZ *mdka* single mutants, respectively. Data are presented as truncated violin plots, where center lines represent the median and limits show the first and third quartiles, respectively. Each scatter dot indicates a C-division event. Statistical analysis was performed using Estimation Stats (https://www.estimationstats.com), between WT and MZ *ptprz1b* or MZ *mdka* with a CI of 99%. (**C**)Quantification of relative angles of C-divisions to mediolateral axis in WT, MZ *ptprz1b* and MZ *mdka* single mutants. Divisions horizontal to mediolateral axis are defined as 0°, while divisions perpendicular to mediolateral axis are defined as 90°. Data were presented as cumulative frequency graph. A Kolmogorov-Smirnov test was performed with a CI of 95% to compare the mean between WT (N = 4) and MZ *ptprz1b* (N = 3) or MZ *mdka* (N = 3). (**D-F**) Representative trajectories of nuclei undergoing C-divisions in WT (**D**), MZ *ptprz1b* (**E**) and MZ *mdka* (**F**) mutants. Time (*t*) is indicated as minutes:seconds before and after cytokinesis. Trajectories and segmented nuclei of parent and daughter cells intercalating into the original half of the neural rod are shown in magenta, and in cyan when starting contralateral intercalation. Green trajectories represent completion of contralateral intercalation. Yellow lines indicate ectopic midlines. Scale bar = 10 μm.

C-divisions in WT zebrafish happen with a stereotypical division orientation (SDO) that is predominantly horizontal relative to the mediolateral axis (Quesada-Hernandez *et al*, 2010). Quantification of planar angles of cell divisions revealed that the C-divisions were not only misplaced but also disorientated in MZ *ptprz1b* and MZ *mdka* mutants prior to midline formation (**Fig 5C**). In WT, C-divisions occurred in predominantly horizontal orientations relative to the mediolateral axis (0-10°) (**Fig 5C; Table 2**). In MZ *ptprz1b* and MZ *mdka* mutants, in contrast, statistically more C-division events took place with diagonal orientations (30-40°, WT vs MZ *ptprz1b, P* < 0.05; WT vs MZ *mdka, P* < 0.05, CI = 95%) (**Fig 5C; Table 2**). These disorientated C-divisions imply that neural progenitor cells in mutant rhombomeres had defective cell polarity.

**Table 2.** Quantification of relative C-division orientation between 30-40°. Data are shown in mean ± SD, Statistical analysis was performed using Estimation Stats (https://www.estimationstats.com), between WT and MZ *ptprz1b* or MZ *mdka* with a CI of 95%.

| Genotype<br>Angles | WT | MZ *ptprz1b* | *P* value | MZ *mdka* | *P* value |
| --- | --- | --- | --- | --- | --- |
| 31-40° | 0.02 ± 0.03 | 0.16 ± 0.09 | *P* <0.0001 | 0.10 ± 0.04 | *P* = 0.0288 |

### Impaired planar cell polarity in MZ *ptprz1b* and MZ *mdka* mutants is rescued by Prickle overexpression

Previous studies had established noncanonical Wnt/Planar cell polarity (PCP) signaling as crucial regulator of midline formation in the zebrafish CNS (Ciruna *et al*., 2006). Mutations in key Wnt/PCP components, such as *van gogh-like 2* (*vangl2*), *wnt5* and *wnt11* resulted in disorientated C-divisions and ectopic midline formation (Ciruna *et al*., 2006). Similar phenotypes were observed in MZ *ptprz1b* and MZ *mdka* mutant hindbrains suggesting impaired cell polarity of neural progenitors also in MZ *ptprz1b* and MZ *mdka* mutants. The fact that midlines were formed, albeit at ectopic positions, suggested that a loss of anteroposterior rather than apicobasal polarity caused the duplicated midlines. In zebrafish, noncanonical Wnt/PCP signaling drives Vangl2-Prickle (Pk) accumulation at the anterior end of neural progenitors to establish polarity (Ciruna *et al*., 2006). A loss of this anteroposterior polarity in *vangl2* mutants is accompanied by Prickle mis-localization (Ciruna *et al*., 2006). To test whether Vangl2-Pk accumulation was disrupted in MZ *ptprz1b* and MZ *mdka* mutants, mRNA encoding a *Drosophila* Prickle-EGFP reporter (Ciruna *et al*., 2006) was injected to visualize endogenous Vangl2 localization. Notably, WT, MZ *ptprz1b* and MZ *mdka* embryos exhibited correct anterior Prickle-EGFP localization in most of the neural progenitors in the hindbrain at 14 to 16 hpf (**Fig 6A-C**). This suggested that endogenous Vangl2 was active and recruited exogenous Prickle-EGFP to the anterior end of neural progenitor cells. We also observed less Prickle puncta in 4 out of 9 MZ *mdka* embryos compared to WT, suggesting a potential reduction of endogenous Vangl2 levels. Importantly, however, overexpression of exogenous *prickle-EGFP* was sufficient to partially rescue the ectopic midline phenotype in MZ *ptprz1b* and MZ *mdka* mutants. 12 out of 14 MZ *ptprz1b* mutants and 5 out of 9 MZ *mdka* mutants injected with *prickle*-*EGFP* mRNA showed formation of a single midline at 16 to 17 hpf (**Fig 6D-G**). This finding thus suggested that ectopic midline formation was caused by a deficiency of endogenous Prickle rather than disruption of endogenous Vangl2 localization. To test for a deficiency in endogenous Prickle, relative qPCR of four annotated zebrafish *prickle* genes, *pk1a, pk1b, pk2a* and *pk2b* was performed. At a pre-symptomatic stage (14 hpf), relative expression levels of *pk1a, pk2a*, and *pk2b* showed no significant difference between WT and MZ *ptprz1b* as well as *mdka* mutants (**Fig 6H**). However, in both mutants, transcription of *pk1b* was significantly downregulated by a factor of 0.62 ± 0.28 (MZ *ptprz1b* mutants) and 0.67 ± 0.17 (MZ *mdka* mutants) (mean ± SD), respectively (**Fig 6H**). Together, our findings suggest that a disruption of Mdka-Ptprz1b signalling results in a deficiency in endogenous Pk1b, thereby disrupting PCP and causing the formation of ectopic midlines. In line with this, overexpression of *Drosophila prickle* rescued the ectopic midline phenotype in MZ *ptprz1b* mutants and MZ *mdka* mutants.

**Figure 6.**
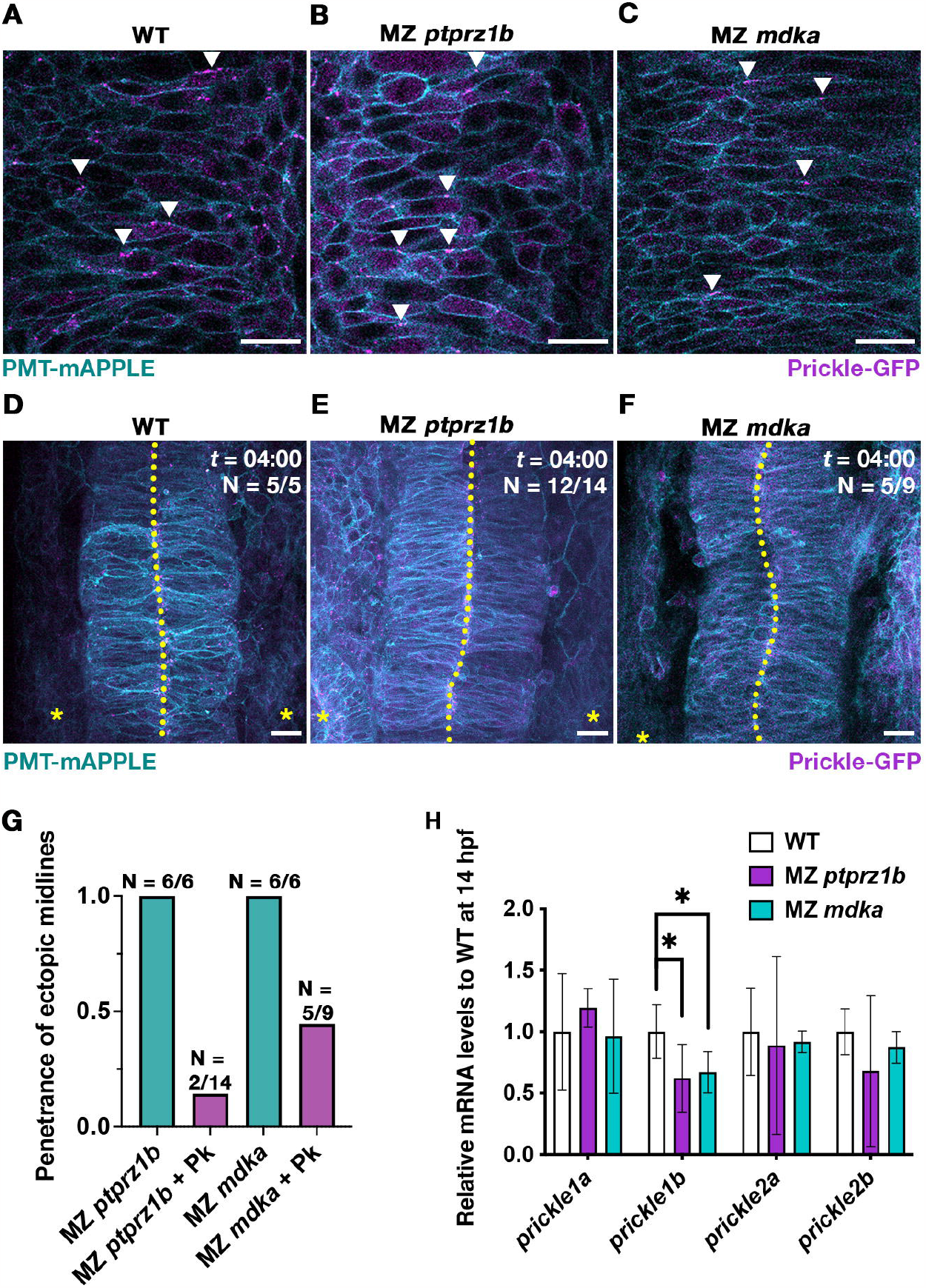
*Drosophila* Prickle-EGFP rescues midline formation in MZ *ptprz1b* and MZ *mdka* mutants that have downregulated *prickle1b* expression. (**A-C**) Representative single-plane images (dorsal views) of neural progenitors in WT (**A**), MZ *ptprz1b* (**B**) and MZ *mdka* mutant rhombomeres (**C**) showing localization of injected *Drosophila* Prickle-EGFP at 14 hpf. Arrowheads indicate anteriorly localized Prickle puncta. Scale bar = 20 μm. (**D-F**) MIP images showing normal midline morphology in rhombomeres at 17 hpf in WT (**D**), MZ *ptprz1b* € and MZ *mdka* single mutants (**F**) injected with *prickle-EGFP* mRNA. Scale bar = 20 μm. (**G**) Quantification of ectopic midline penetrance in MZ *ptprz1b* and MZ *mdka* mutants injected with *PMT-mEGFP*/*PMT-mApple* mRNA with and without *prickle-EGFP* mRNA. Data for *PMT-mEGFP*/*PMT-mApple* mRNA injected mutants were taken from (**Figs. 4** and **5**). Sample numbers are indicated on top of each bar. (**H**) Relative qPCR quantification of *prickle1a, prickle1b, prickle2a* and *prickle2b* expression at 14 hpf.

## DISCUSSION

Midline formation in the CNS is crucial for mirror-symmetric distribution of neural progenitor cells and later neural circuits (Tawk *et al*., 2007). The noncanonical Wnt/PCP signalling pathway, acting over long-range through noncanonical Wnt ligands, is a major determinant in midline formation, and acts through cell convergence/extension and polarized C-divisions in the zebrafish CNS (Ciruna *et al*., 2006; Quesada-Hernandez *et al*., 2010). This pathway orchestrates cell convergence/extension and polarized C-divisions during midline formation in the zebrafish CNS (Ciruna *et al*., 2006; Quesada-Hernandez *et al*., 2010). However, it remains poorly understood whether and how other extrinsic cues synergize with or even coordinate assembly of noncanonical Wnt/PCP components (Hirano *et al*., 2022). In this study, we provide evidence that the zebrafish heparin-binding growth factor Mdka acts up-stream of the noncanonical Wnt/PCP pathway. We identified a so far undescribed role for Mdka/Ptn-Ptprz1b signaling in neural keel convergence and contralateral cell intercalations after midline-crossing cell divisions (C-divisions) that are needed to facilitate midline formation in the hindbrain. Rescue experiments placed Mdka-Ptprz1b upstream of noncanonical Wnt/PCP signaling suggesting possible crosstalk between both pathways. Our findings indicate that Mdka-Ptprz1b signaling is a new extrinsic cue that regulates hindbrain planar cell polarity upstream of Wnt/PCP signaling.

### Maternal Ptprz1b and zygotic Mdka control midline formation

Our mutant analysis identified defective midline formation in rhombomeres of zebrafish embryos deficient for Mdka, Ptn and Ptprz1b. In contrast to WT, MZ *mdka*, MZ *ptn* and MZ *ptprz1b* mutants formed duplicated midlines at ectopic positions in the hindbrain during neurulation at 16 to 17 hpf. MZ *mdka* and MZ *ptprz1b* mutants exhibited similar phenotypic penetrance affecting r1, r2, r4 and r5, while MZ *ptn* and MZ *ptprz1b* mutants showed similar penetrance in r5 to r7. The similarity of hindbrain phenotypes in ligand and receptor mutants suggested a possible interaction between Mdka and Ptn with Ptprz1b, respectively. Such an interaction was confirmed by PLA and FCCS *in vivo*. Together, this suggests that diffusible Mdka and Ptn, which originate from distinct rhombomeres, bind to Ptprz1b with different affinities to coordinate midline formation during dynamic hindbrain morphogenesis.

Importantly, at 14 hpf, levels of endogenous zygotic *mdka* but not of *ptprz1b* expression were elevated in those rhombomeres that manifested ectopic midlines in MZ *mdka* and MZ *ptprz1b* mutants. In addition, depletion of maternal *ptprz1b* by crossing MZ *ptprz1b* females with WT males was sufficient to induce aberrant midlines in r2 and r3, while embryos from the reverse cross showed normal midline formation. This indicated that a loss of maternal rather than zygotic Ptprz1b affects midline formation. In contrast, the loss of maternal *mdka* had no observable effect on midline morphology, suggesting that it is zygotic *mdka* expression that regulates midline formation. This is consistent with a high abundance of maternal *ptprz1b* mRNA, especially when compared to *mdka*. Together, this implies that at neurulation stages, zygotic Mdka ligand, locally produced in r1 to r5, diffuses across rhombomeres to interact with maternally provided Ptprz1 receptor that is broadly distributed in neural progenitors. The degree of Mdka-Ptprz1b signaling thus seems to be defined by the restricted spatiotemporal pattern of *mdka* expression.

Using Ptn as an example, we showed that ligand binding effectively induced internalization of Ptprz1b. Such internalization may prevent repetitive activation of Ptprz1b signaling. In addition, it ultimately reshaped the originally uniform receptor distribution *in vivo*. This observation suggested that gradients of zygotically expressed Mdk and Ptn ligands can modulate distribution of maternally provided receptors so that levels of signaling activation are precisely controlled in order to coordinate cell convergence and division in an accurate spatiotemporal pattern. We demonstrated that this is consistent with a reaction-diffusion model, in the formation of opposing gradients of ligand and receptor expression. When the receptor expression was increased in simulation, we observed a decrease in the decay length of the ligand, flattening the ligand’s concentration profile (**Fig S6**).

Using PLA *in vivo*, we found that Ptn exhibited a considerably higher affinity to Ptprz1b (mean ± SD = 0.36 ± 0.13) than Mdka (mean ± SD = 0.17 ± 0.05) under the same molar ratios. However, MZ *ptn* mutants showed an overall milder severity and lower penetrance of midline phenotypes when compared to MZ *mdka*. This was probably due to the restricted zygotic expression of *ptn*, which at 14 hpf is limited to r5 and r6. Despite co-expression of *mdka* and *ptn* in these two rhombomeres, mutant phenotypes were observed in both single mutants. This suggested that *ptn*, despite having higher binding affinities, could not compensate for the loss of *mdka* in r5 and r6 as well as more anterior rhombomeres. On the other hand, even high *mdka* expression levels were insufficient to fully compensate for the loss of *ptn*. In agreement with a partial compensation by *mdka*, the penetrance of the midline phenotype was much lower in MZ *ptn* mutants when compared to either MZ *ptprz1b* or MZ *mdka* mutants. Together, these data suggest a complex interplay of Mdk/Ptn ligands that are structurally related but differentially expressed in rhombomeres and exhibit distinct affinities to their Ptprz1 receptors. These ligands set up a pattern in the hindbrain that triggers receptor internalization, thus shapes the distribution of membrane-bound receptor and thereby precisely controls signaling. In the context of compensation, while all ligands are likely to have at least partially overlapping roles, they appear to be indispensable in regulating rhombomeric midline formation. The targets of Ptprz1b signaling in the hindbrain are so far unknown. It will be interesting to see in the future how such targets are regulated by an interplay of related Mdk/Ptn ligands to control coordinated cell behavior during rhombomere convergence and midline formation. Interestingly, it is also a combination of Wnt4, Wnt5 and Wnt11 ligands that controls noncanonical PCP signaling during neural plate convergence (Ciruna et al., 2006).

Importantly, Mdkb exhibited significant binding affinity to Ptprz1b (mean ± SD = 0.17 ± 0.09) and is broadly expressed in r1 to r6, similar to *mdka*. However, *mdkb* failed to compensate for midline formation in MZ *mdka* mutants. As *mdka* and *mdkb* are largely co-expressed in the same rhombomeres, one possible explanation is that Mdkb binding to Ptprz1b triggers distinct downstream pathways that are not related to midline formation in rhombomeres. Consistent with this, we did not observe ectopic midlines in MZ *mdkb* mutants (**Fig S4A, B, A’-B’**). Instead, midline formation was significantly delayed in MZ *mdkb* mutants, and a distinctive midline could only be observed at 18 hpf (**Fig S4C-D, C’-D’**). These phenotypic differences in *mdkb* vs. *mdka*/*ptn* mutants are consistent with the idea of a functional divergence of midkine ligands involving distinct downstream pathways, which has been proposed previously (Calinescu *et al*, 2009; Winkler *et al*., 2003).

### Mdka-Ptprz1b signaling acts upstream of the PCP component Prickle

The ectopic midlines observed in MZ *mdka*, MZ *ptn* and MZ *ptprz1b* mutants were phenotypically very similar to those reported for several Wnt/PCP mutants, including MZ *van gogh-like 2* (*vangl2*)/*trilobite* (*tri*) and MZ *wnt11*/*silberblick* (*slb*); MZ *wnt5*/*pipetail* (*ppt*) double mutants. These mutants also exhibit duplicated mirror-symmetric ectopic midlines formed in the hindbrain and spinal cord, caused by a loss of anteroposterior polarity in neural progenitors (Ciruna *et al*., 2006; Tawk *et al*., 2007). This loss was reported to lead to defective neural keel convergence to place neural progenitor cells away from the middle of the neural keel (Tawk *et al*., 2007). Consequently, cell division occurred at ectopic positions lateral to the presumptive midline, and the daughter cells subsequently failed to intercalate into the contralateral half of the neural keel (Ciruna *et al*., 2006; Tawk *et al*., 2007). The aberrantly positioned progenitors still established apicobasal polarity and remained in register but aligned their apical ends at an ectopic plane, which resulted in duplicated midlines with disorganized cells in between them (Tawk *et al*., 2007). Similarly, MZ *mdka* and MZ *ptprz1b* mutant neural keels showed a significant delay in convergence that was particularly evident in rhombomeres r2, r3 and r4. In WT, r2 and r3 are broader than other rhombomeres at onset of convergence and always appeared as most severely affected in mutants. We thus speculate that Mdka-Ptprz1b signaling could be important to facilitate accelerated convergence of r2 and r3 so that consequently all seven rhombomeres have comparable width to allow C-divisions at aligned medial positions. Importantly, MZ *mdkb* mutants exhibited a narrower r2 than WT suggestive for a possible acceleration of convergence and exhibited delayed midline formation rather than ectopic midlines (**Fig S4G-I**). This supports our speculation that ectopic midlines are a consequence of delayed convergence in rhombomeres. We thus speculate that a convergence delay in MZ *mdka* and MZ *ptprz1b* mutants led to ectopic positioning of C-divisions and a failure of progenitors to contralaterally intercalate (**Fig 7A**).

**Figure 7.**
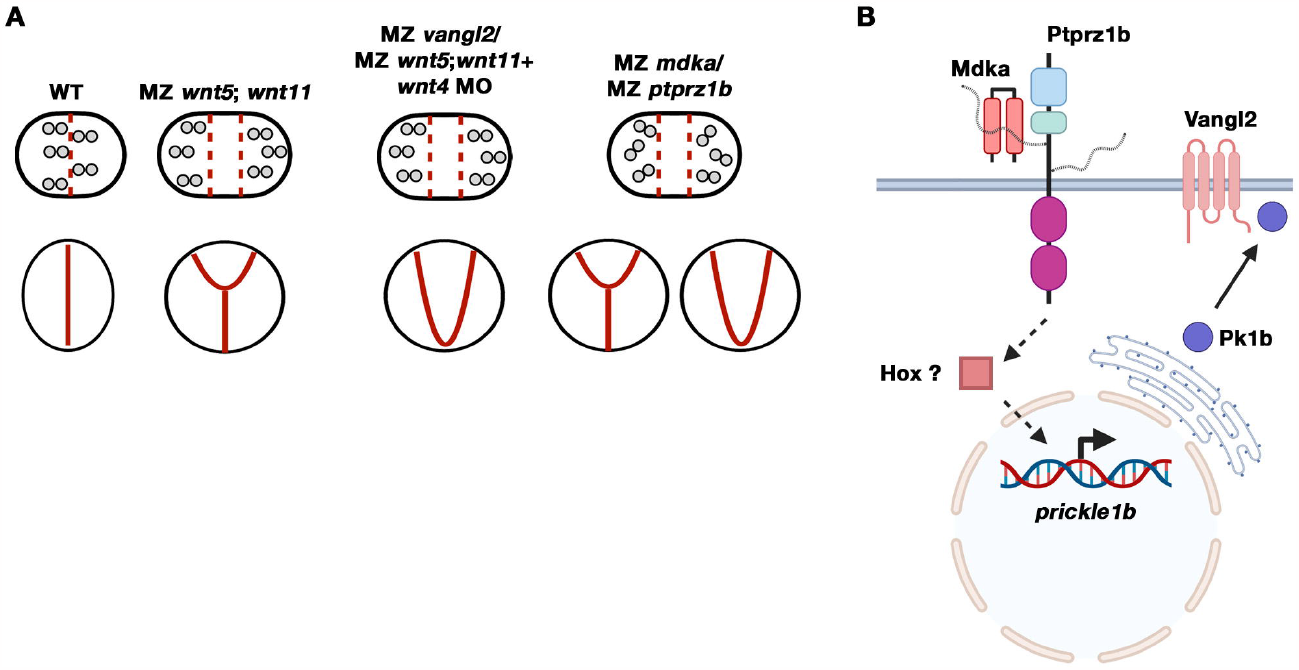
Crosstalk of Mdka-Ptprz1b with noncanonical Wnt/PCP signalling through *prickle1b*. (**A**)Illustration of convergent extension (CE), stereotypical cell-division orientation (SDO) and ectopic midline phenotypes in different zebrafish mutants or morphants. Upper panel shows schematic dorsal views of individual rhombomeres at 15 hpf. Width of rhombomeres indicates extent of convergence, and dotted lines represent presumptive midlines where C-divisions take place. Lower panel shows schematic transverse views of corresponding rhombomeres at 17 hpf. Solid lines reflect midline morphologies. Noncanonical Wnt/PCP mutants, MZ *mdka* and MZ *ptprz1b* mutants exhibit limited convergence and ectopic C-divisions at lateral positions. SDO was additionally altered in MZ *mdka* and MZ *ptprz1b* mutants. (**B**)Hypothetical model of signalling crosstalk between Mdka-Ptprz1b and noncanonical Wnt/PCP pathway. Figure is created with BioRender.com
.

The striking similarities in mutant midline phenotypes suggested a possible crosstalk between Mdka-Ptprz1b and noncanonical Wnt/PCP pathways. At 14 hpf, neural progenitor cells in the spinal cord exhibit anterior Vangl2-Prickle (Pk) localization at the plasma membrane (Ciruna *et al*., 2006). At this stage, both MZ *mdka* and MZ *ptprz1b* mutants had correctly localized Pk near the anterior plasma membrane of neural progenitors, suggesting that localization and function of Vangl2 were normal in both mutants. Unexpectedly, however, we also observed that overexpression of *Drosophila* Prickle partially rescued the ectopic midline phenotypes in MZ *mdka* and MZ *ptprz1b* mutants, suggesting a deficiency of endogenous Prickle as cause for ectopic midline formation. Consistent with the latter, previous studies had reported that a knockdown of the zebrafish prickle ortholog *pk1a* induced ectopic midlines in the hindbrain (Tawk *et al*., 2007). Consistent with this hypothesis, we found that transcription of the co-ortholog *pk1b*, but not *pk1a*, was significantly downregulated in MZ *mdka* and MZ *ptprz1b* mutants. These data strongly suggest that Mdka-Ptprz1b acts upstream of noncanonical Wnt/PCP signaling and is involved in regulating transcription of *pk1b* at correct times and places (**Fig 7B**). Future studies need to identify transcriptional regulators downstream of Mdka/Ptprz1b that control transcription of *pk1b*.

It has been reported that at 16 hpf, *pk1b* is expressed in r4 and weakly in cells at the lateral edges of more anterior rhombomeres (Rohrschneider *et al*, 2007). Its expression is regulated by Hoxb1a, which in turn is stimulated by retinoid acid (RA) secreted from somites positioned immediately adjacent to the hindbrain (Rohrschneider *et al*., 2007; Weicksel *et al*, 2014). Midkine genes are also known to be RA-inducible (Matsubara *et al*, 1994; Tomomura *et al*, 1990; Winkler & Moon, 2001a). Future studies thus need to test whether RA coordinates expression of *mdka*/*ptn* and *hox* genes in the hindbrain. Such an interaction could then lead to spatiotemporal control of *prickle* transcription to establish a regionally restricted distribution of PCP components in rhombomeres.

Apart from delayed convergence, we also observed disorientated C-division in both *mdka* and *ptprz1b* mutants, which strongly resembles previously reported phenotypes in MZ *fzd7a*; MZ *fzd7b* double mutant (Quesada-Hernandez *et al*., 2010). However, *mdka* and *ptprz1b* mutants form duplicated midlines rather than no midlines as seen in MZ *fzd7a*; MZ *fzd7b* double mutant. This could be explained by differences in SDO patterning: in *mdka* or *ptprz1b* mutants, disorientated C-divisions occurred in a range between 30-40° relative to the mediolateral axis, while in MZ *fzd7a*; MZ *fzd7b* double mutants, C-divisions were reported in a range between 30-90°.

Another proposed target of PTPRZ1b is Rho/ROCK signaling (Bhaduri *et al*., 2020; Qin *et al*., 2017). Interestingly in zebrafish, noncanonical Wnt/PCP signaling has been shown to control Rho/ROCK activity to facilitate convergence and extension during gastrulation (Bai *et al*, 2014; Marlow *et al*, 2002; Zhu *et al*, 2006). Future studies thus need to examine whether Mdk/Ptn-Ptprz1 controls neural keel convergence by regulating Rho/ROCK, and whether such a regulation is mediated through the noncanonical Wnt/PCP pathway.

In conclusion, we propose a model where maternally provided Ptprz1b receptor is activated by locally produced Mdka or Ptn ligands to control transcription of *pk1b*. This in turn sets up a localized distribution of PCP pathway components that is required to correctly position C-divisions during hindbrain morphogenesis. In this process, distribution of Ptprz1b is dynamically reshaped by zygotic Mdka expression, which controls receptor internalization and thus availability of signaling receptor on cell membranes. Hence, regionally restricted availability of Mdka ligand, in combination (or competition) with high-affinity Ptn, and possibly also Mdkb, determines the level of Ptprz1b activation to control noncanonical PCP in neural progenitors. Together, this modulates the speed of cell convergence that is needed to correctly time and place C-divisions in the forming rhombomeres.

## MATERIALS AND METHODS

### Reagents and tools table

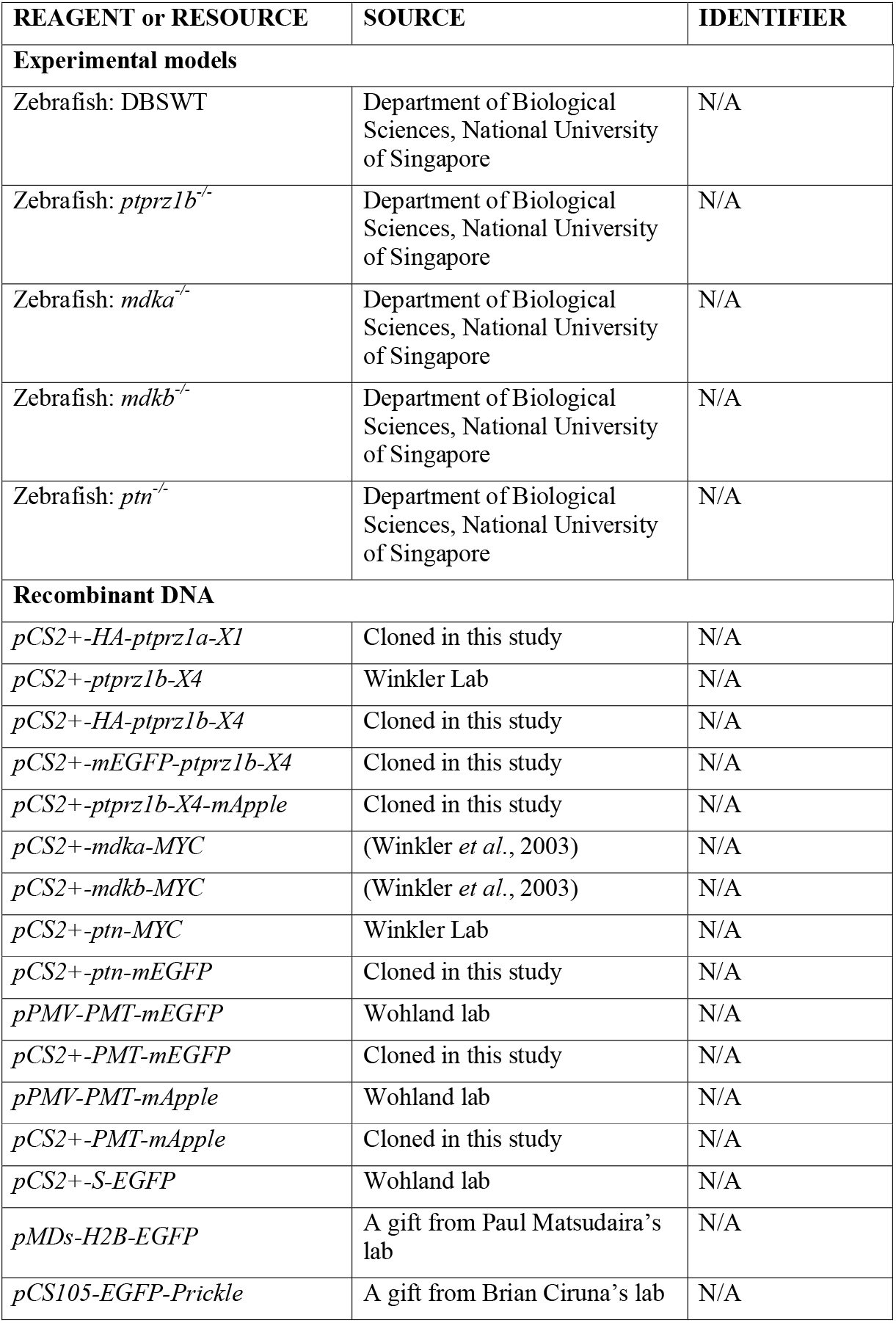
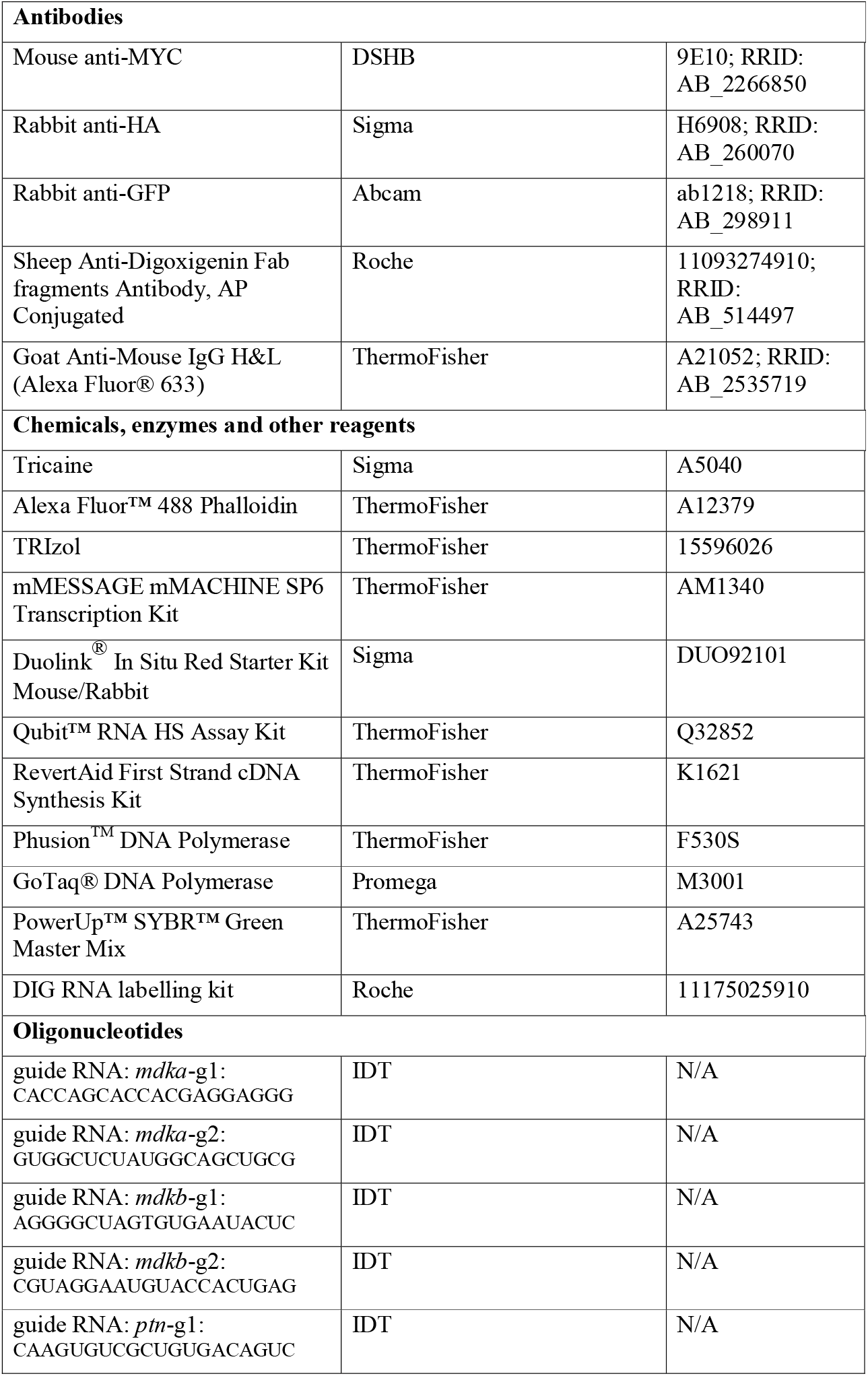
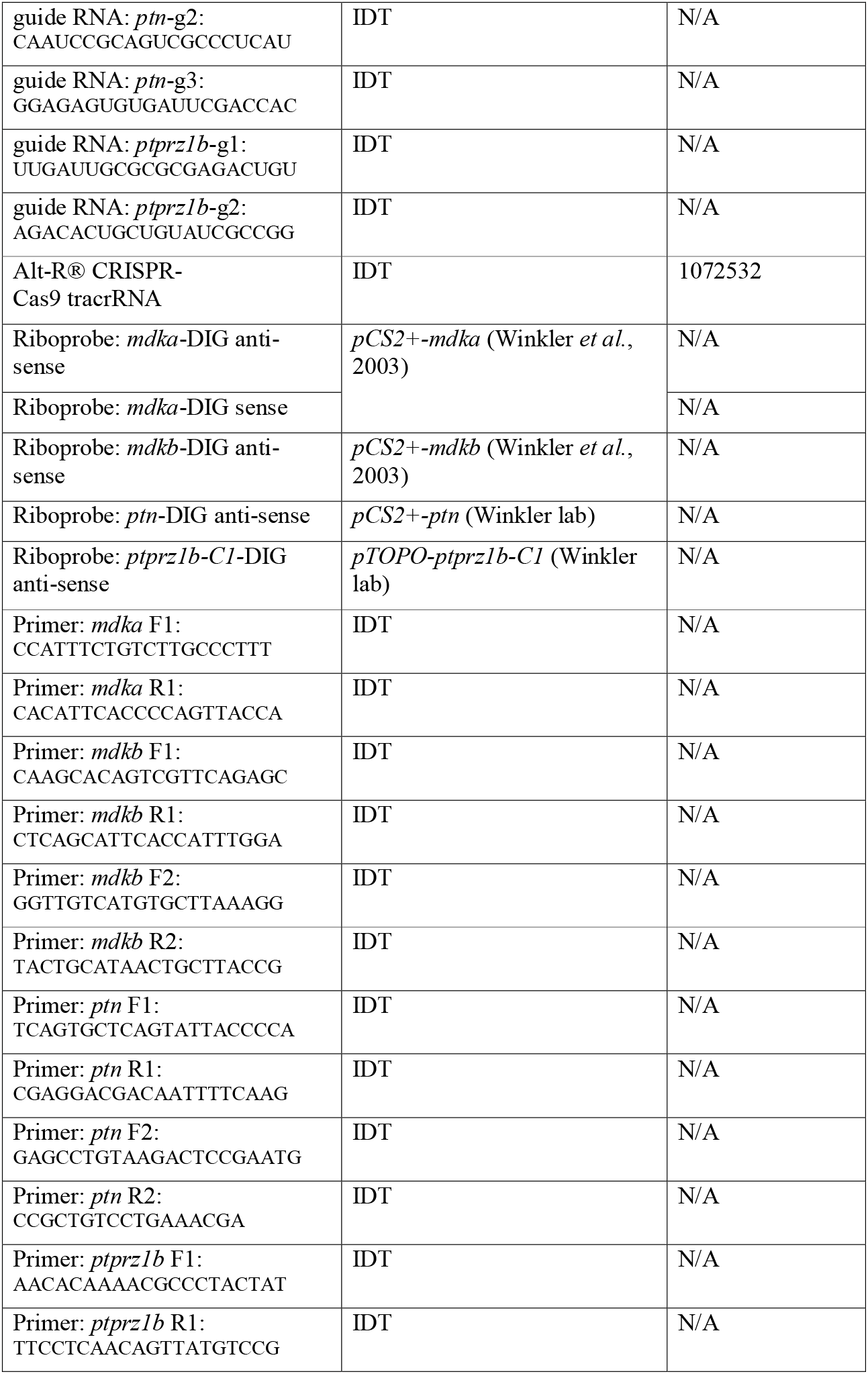
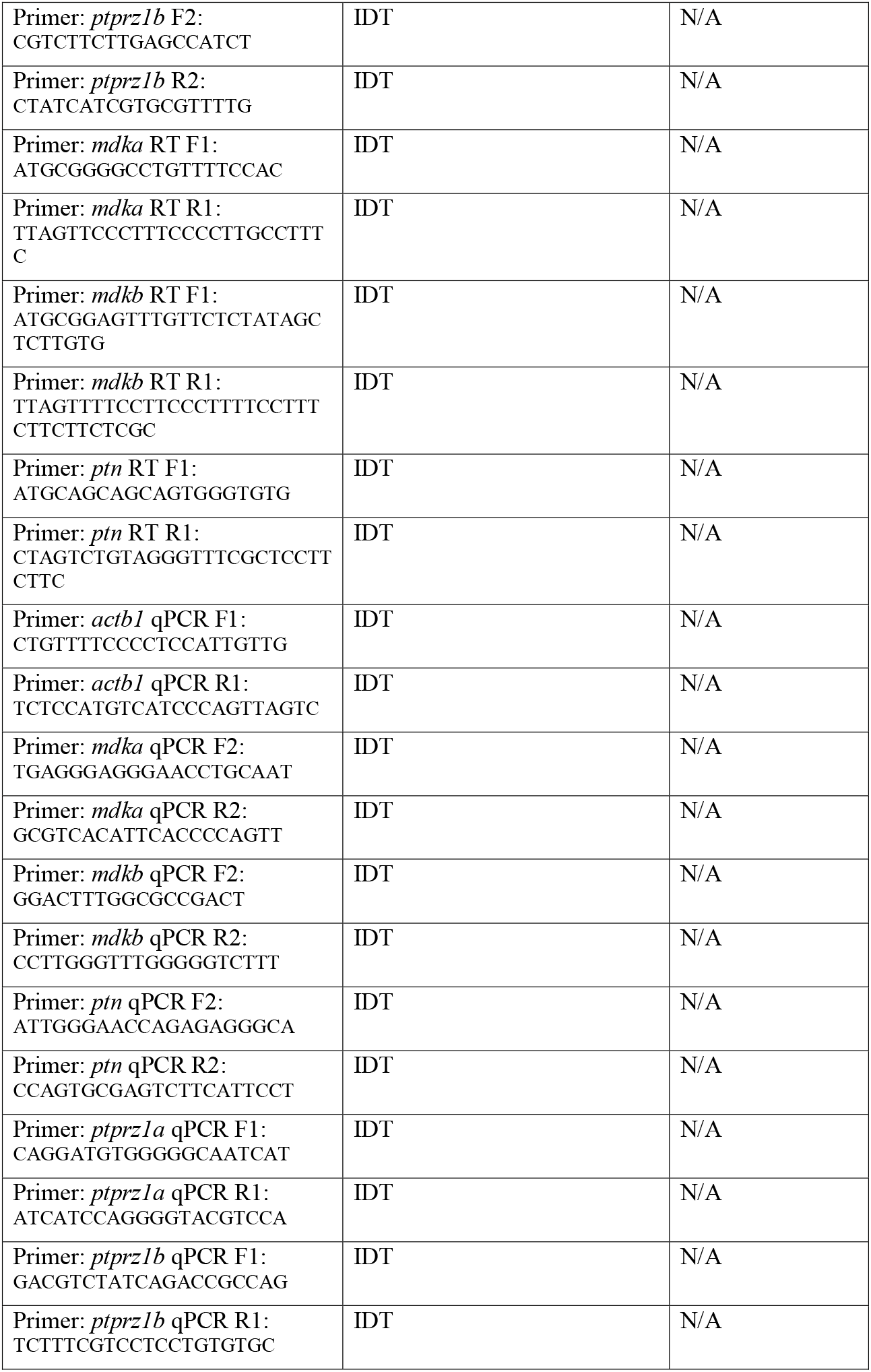
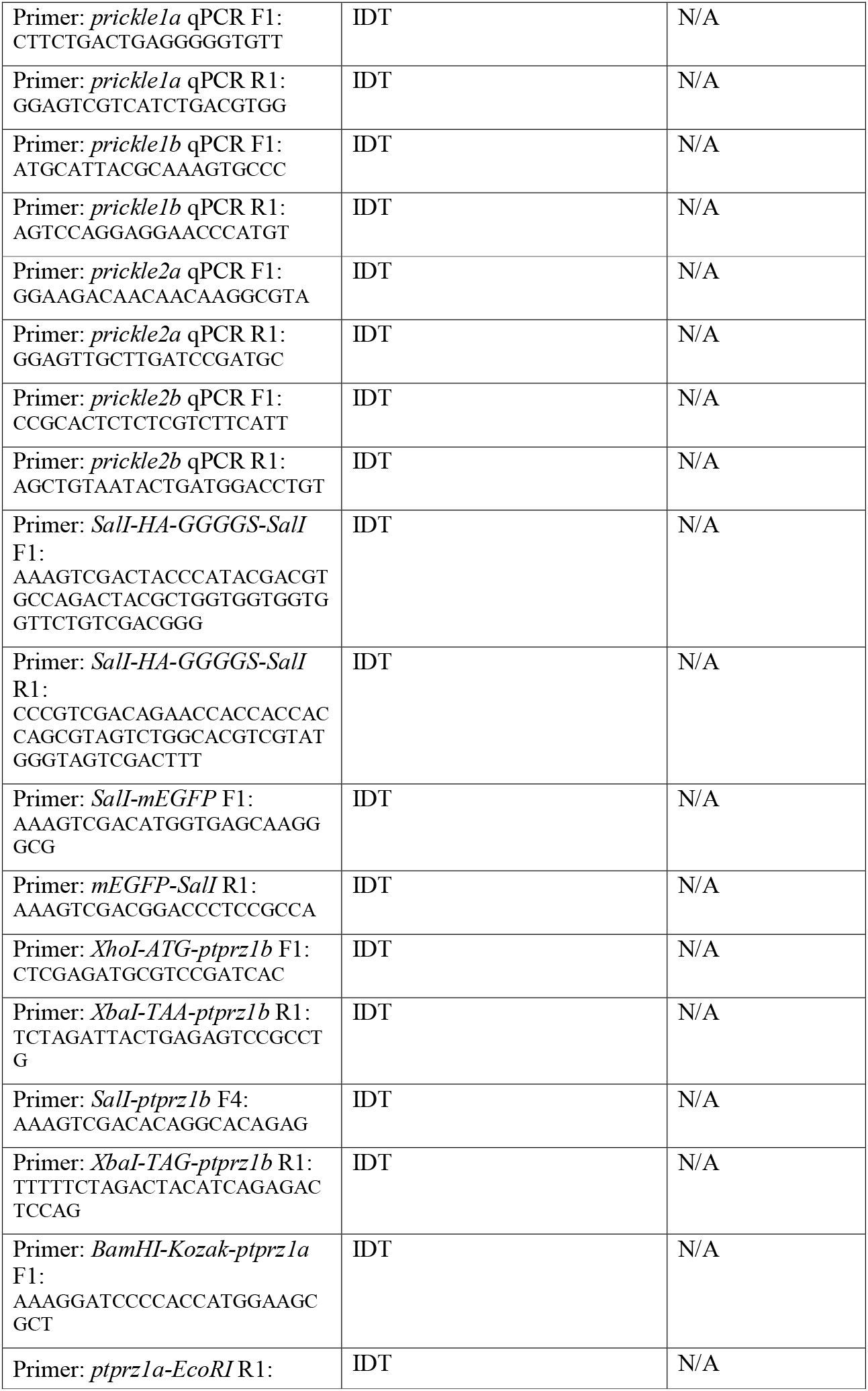
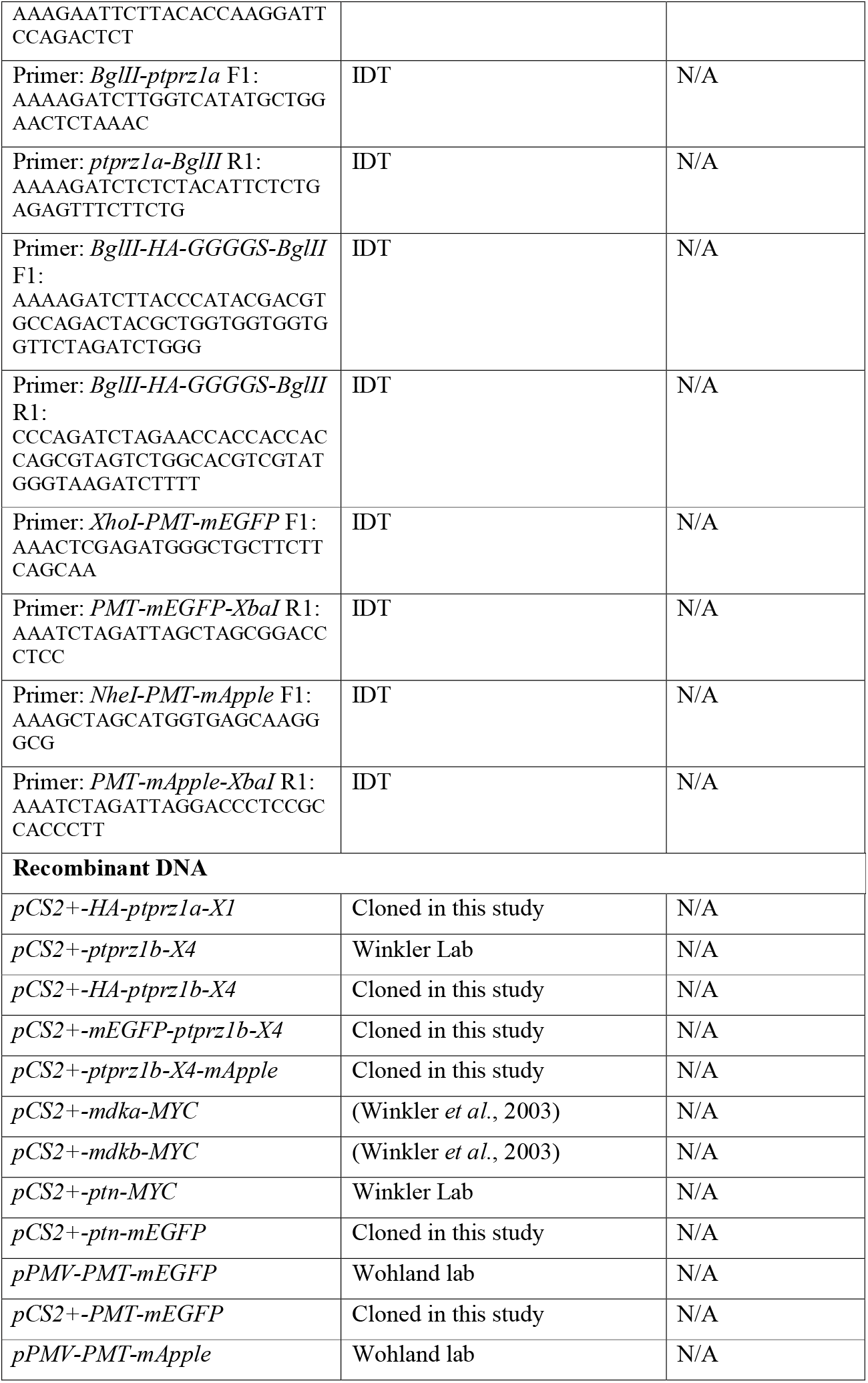
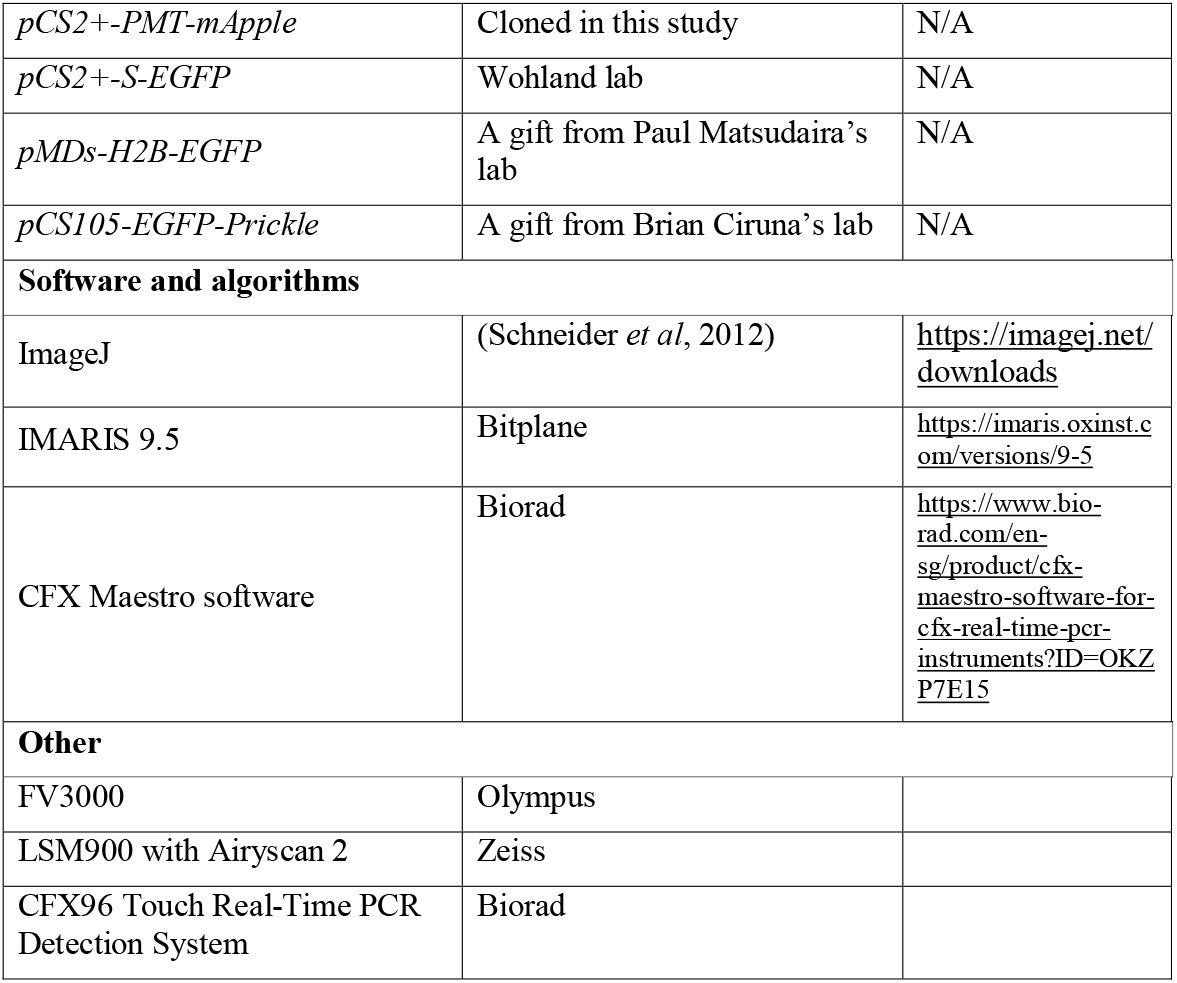

## Method and protocols

### Establishment of zebrafish mutants

All animal experiments were conducted in accordance with protocols (BR15-0119, BR19-0120, BR22-1497) approved by the Institutional Animal Care and Use Committee (IACUC) of the National University of Singapore. Adult zebrafish were housed in 28□ circulating systems under a 14h/10h light/dark cycle in the fish facility of the Department of Biological Sciences (DBS) at National University of Singapore. Wild type (WT) and mutant zebrafish embryos were obtained by crossing corresponding adult male and female fish. Embryos under 5 days post fertilization (dpf) were cultured in 0.3X Danieau’s solution (17.4 mM NaCl, 0.21 mM KCl, 0.12 mM MgSO_4_, 0.18 mM Ca(NO_3_)_2_, 1.5 mM HEPES, pH = 7.2) in a 28□ incubator. Embryonic stages were defined by hours post fertilization (hpf) at 28□ or by morphological features (Kimmel *et al*, 1995). Zebrafish larvae older than 5 dpf were raised in static tanks with regular changes of water for a maximum of 3 weeks before shifting into a circulating system.

All zebrafish mutants were generated by CRISPR-Cas9 gene editing. CRISPR RNAs (crRNA) were designed using CCTop (https://cctop.cos.uni-heidelberg.de:8043) or the IDT online prediction application (https://sg.idtdna.com). All crRNAs and tracrRNAs were synthesized by IDT (Singapore). CRISPR-Cas9 ribonucleoprotein (RNP; IDT Singapore) assembly was performed according to the manufacturer’s protocol with minor modification. Briefly, 3 μL of crRNAs (100 μM, IDT) and tracrRNAs (100 μM, IDT) were added to 94 μL of Nuclease-Free Duplex Buffer (IDT), subsequently heated at 95□ for 5 min and chilled on ice to allow annealing into guide RNAs (gRNA). 3 μL of gRNAs were then mixed with 3 μL of Cas9 working solution (0.5 μg/μL, IDT) with subsequent incubation at 37□ for 10 min to form RNP complexes. Approximately 3 nL of RNP solution were injected into 1-cell stage embryos collected from DBS wildtype (WT) incrosses. Injected embryos were raised to 2 months post fertilization (mpf) and then fin clipped for genotyping by PCR with respective primers. PCR positive individuals were regarded as potential F0 founders and outcrossed with wildtype to produce F1 generations. F1 fish were later genotyped and sequenced to identify the mutated allele. Fish with the same allele were incrossed to produce F2 progeny and establish stable mutant lines. F2 homozygotes were identified by genotyping and further incrossed to generate maternal-zygotic (MZ) mutants.

### DNA construct assembly and mRNA synthesis

For proximity ligation assays (PLA), *pCS2+-HA-ptprz1a-X1* and *pCS2+-HA-ptprz1b-X4* constructs were cloned. The *pCS2+-HA-ptprz1a-X1* was assembled from the predicted sequence of *ptprz1a* splicing variant X1 (*ptprz1a-X1*, NCBI Reference Sequence: XM_005163137.4) with a HA-tag sequence inserted into the N-terminus sequence of *ptprz1a*. The *pCS2+-HA-ptprz1b-X4* carried the predicted sequence of *ptprz1b* splicing variant X4 (*ptprz1b-X4*, NCBI Reference Sequence: XM_021475402.1) and a HA-tag sequence at the N-terminus sequence of *ptprz1b*. To visualize Ptprz1b distribution, *pCS2+-mEGFP-ptprz1b-X4* was subcloned from *pCS2+-HA-ptprz1b-X4* with *HA* sequence substituted by *mEGFP* sequence. Capped mRNA was synthesized using mMACHINE SP6 Transcription Kit (ThermoFisher) with *NotI*-linearized *pCS2+-mdka-MYC, pCS2+-mdkb-MYC, pCS2+-ptn-MYC, pCS2+-HA-ptprz1a-X1, pCS2+-HA-ptprz1b-X4, pCS2+-mEGFP-ptprz1b-X4* and *pCS2*+-*secreted*-*GFP* plasmids as templates, respectively.

For FCCS and FCS measurements, *ptn-mEGFP, ptprz1b-mApple* and *PMT-mEGFP-mApple* mRNAs were synthesized by SP6 Transcription Kit (ThermoFisher) with *MfeI*-linearized *pCS2+-ptn-mEGFP, pCS2+-ptprz1b-mApple* and *pCS2+-PMT-mEGFP-mApple* plasmids as templates.

For confocal time-lapse imaging, *NotI*-linearized *pMDs-H2B-EGFP, MfeI*-linearized *pCS2+-PMT-mEGFP* or *pCS2+-PMT-mApple* and *KpnI*-linearized *pCS105-EGFP-Prickle* plasmids were used as templates for capped mRNA synthesis using mMESSAGE mMACHINE SP6 Transcription Kit (ThermoFisher). The concentration of capped mRNA was determined using Qubit™ RNA HS Assay Kit (Q32852, ThermoFisher).

### Immunohistochemistry

Immunostaining of zebrafish embryos was performed following protocols reported previously (Yao *et al*, 2013). Briefly, samples were fixed with 4% PFA/PBST at 4□ overnight. After a series of PBST washes, permeabilization was performed with 5 μg/mL of proteinase K in PBST for 5 min. After a wash with 2 mg/mL of glycine, samples were re-fixed in 4% PFA/PBST for 20 min at room temperature and subsequently washed with 1X PBST for 5 times, 5 min each. Blocking was achieved by using 5% sheep serum/PBST at room temperature for 1 h. Primary antibodies were diluted in 5% sheep serum/PBST. For MYC staining, mouse anti-MYC, 9E10 (DSHB) was diluted 1:100 in 5% sheep serum/PBST. After removal of primary antibodies, samples were washed with 1X PBST for 5 times, 5 min each. Secondary antibodies, Goat Anti-Mouse IgG H&L (Alexa Fluor® 633) (A21052, ThermoFisher) was applied with a dilution factor of 1:1000 in 5% sheep serum/PBST and incubated overnight at 4[. Excess antibodies were removed by three washes with 1X PBST for 5 min each. After a counterstain with 5 μg/mL DAPI for 10 min, samples were mounted on slides or imaging dishes with Mowiol ® 4-88 for confocal imaging.

### Proximity ligation assay

For proximity ligation assays, *mdka-MYC*/*mdkb-MYC*/*ptn*-*MYC* and *HA*-*ptprz1a*/*HA-ptprz1b* mRNAs were co-injected into WT embryos. The concentration of each component for injection was adjusted so that all ligand-receptor combinations had comparable molar ratios. The detailed concentrations are listed as follow: *mdka-MYC*, 20 ng/μL; *mdkb-MYC*, 20 ng/μL; *ptn-MYC*, 20.7 ng/μL; *HA-ptprz1a*, 41.5 ng/μL; *HA-ptprz1b*, 30 ng/μL. The injection amount was adjusted to 1 nL. Embryos injected with *HA-ptprz1b* mRNA served as negative controls. As positive control, embryos were injected with *mdka-MYC* mRNA and treated with secondary antibodies directed against both (+) and (-) oligonucleotides to recognize anti-MYC primary antibodies. A random collision control was included using embryos injected with 20 pg *mdka-MYC* and 5.6 pg *secreted-GFP* mRNA. Injected embryos were incubated in a 28□ incubator and fixed with 4% PFA/PBST at 70%-80% epiboly. After overnight fixation at 4□, embryos were washed, permeabilized and blocked as described for immunostaining. Primary antibodies, mouse anti-MYC, 9E10, (DSHB) and rabbit anti-HA (H6908, Sigma) or rabbit anti-GFP (ab1218, Abcam) for random collision control, were diluted 1:100 in Duolink® Antibody Diluent provided in the DuoLink In Situ Red Starter Kit Mouse/Rabbit (DUO92101, Sigma), and added to samples for overnight incubation at 4[. Unbound antibodies were washed off by three washes for 5 min each with 1X Wash Buffer A (DUO92101, Sigma). As secondary antibodies, a pair of antibodies labelled with an oligonucleotide barcode, goat anti-mouse MINUS and goat anti-rabbit PLUS (DUO92101, Sigma), were used. Samples were incubated in a humid chamber at 37□ for 1h followed by three washes with 1X Wash Buffer A for 5 min each. To create a template for signal amplification, barcodes were ligated by ligase supplied in the kit at 37□ for 30 min with two subsequent washes with 1X Wash Buffer A for 5 min each. Using the ligated barcodes as templates, signal amplification was performed with a polymerase supplied by the kit at 37□ for 100 min. 1X Wash Buffer B (DUO92101, Sigma) was used to remove excess fluorophore-tagged probes by washing for 10 min twice. Washed embryos were then transferred to 0.01X Wash Buffer B and mounted with Mounting Medium with DAPI (DUO82040, Sigma) on an imaging dish for confocal imaging on FV3000 (Olympus).

In order to create two distinct groups of cells that expressed Ptn-MYC and mEGFP-Ptprz1b, respectively, at dome stage, 8-cell stage injection was performed. 40 pg of *mEGFP-ptprz1b* mRNA was injected into one of the cells. A mixture of 8 pg of PMT-mEGFP and 40 pg *ptn-MYC* mRNA was injected into another distally located cell. Injected embryos were fixed at dome stage for immunostaining or PLA using mouse anti-MYC, 9E10, (DSHB) and rabbit anti-GFP (ab1218, Abcam).

To create a confined source of Ptn-MYC and homogenous distribution of mEGFP-Ptprz1b or PMT-mEGFP, embryos were injected at 1- and 8-cell stages, respectively. At 1-cell stage, 40 pg *mEGFP-ptprz1b* or 8 pg *PMT-mEGFP* mRNA was injected to allow ubiquitous and uniform distribution of the expressed protein. At 8-cell stage, 20 pg *ptn-MYC* mRNA was introduced into a single cell to create a domain of cells expressing Ptn-MYC at later stage. Injected embryos were incubated at 28□ before fixation at dome stage. To indicate the source of Ptn in 1- and 8-cell stage injected embryos, samples were immunostained with mouse anti-MYC, 9E10, (DSHB) and subsequently Goat AntiMouse IgG H&L (Alexa Fluor® 633) (A21052, ThermoFisher; 1:1000) as described above. After washing off unbound antibodies, samples were mounted with Mounting Medium with DAPI (DUO82040, Sigma) and subjected to confocal imaging on LSM900 (Zeiss).

## FCS and FCCS measurement

FCS and dual colour-FCCS (DC-FCCS) measurements were conducted on the animal pole of 4 hpf zebrafish embryos following published protocols with modifications (Foo *et al*, 2011; Ma *et al*, 2014). Before measurement, the respective structure parameter (κ), diffusion time (τ_d_) and effective observation volume (V_eff_) of Atto 488 and Atto 565 dye was acquired for calibration. For FCS, Atto 488 dye was excited under 485 (8.5 μW) pulsed laser and for FCCS, Atto 488 and Atto 565 dyes were simultaneously excited by 488 (6 μW) and 543 nm (8-10 μW) continuous-wave lasers, respectively. Fluorescence fluctuations from dye and EGFP were recorded and subsequently converted into auto-correlation functions (ACF) for FCS, recorded fluorescence fluctuations from dye, EGFP and mApple were converted into both ACF and a cross-correlation function (CCF) for FCCS by the software SymPhoTime 64 (PicoQuant, Germany). ACFs and CCFs were imported into Igor Pro 8 (Wavemetrics, USA) for curve fitting by a 3D-2 particles (3D2P1t) model in a self-written programme (https://www.dbs.nus.edu.sg/lab/BFL/confocal_FCS.html).

For FCS, the ACF is expressed as:

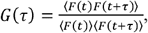

where *F*(*t*) is the fluorescence intensity at time *t, F*(*t +τ*) is the fluorescence intensity a time τ later, and ⟨ … ⟩ denotes time averaging.

*G*(τ) for a three-dimensional free diffusion process with two components and triplet state (3D2P1t) is given by:

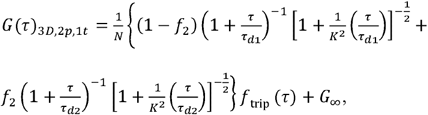

where *f*_2_ is the fraction of second component, with the first fraction being *f*_1_=1 − *f*_2_

For FCCS, the ACF of the green channel (G) is expressed as:

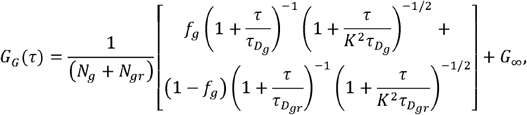

while ACF of red channel (R) is expressed as:

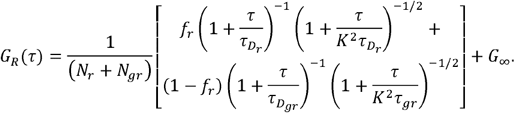

The cross-correlation function is written as:

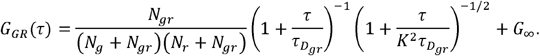

In DC-FCCS, the exact amount of cross-correlation could not be quantitated due to spectral bleed-through. Hence, the relative cross-correlation(Q) was introduced to compare the percentage of cross-correlation of samples with that of negative and positive controls, respectively. In theory, Q is the ratio of the amplitudes of the CCF to the individual ACFs and is calculated as:

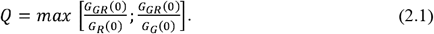

From equation (2.1), the amplitude of ACF (G(0) is inversely proportional to the number of molecules (N). Since N was derived as a parameter from fitting, the equation for Q was re-written in terms of N as:

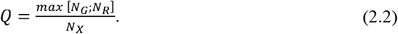

To determine the range of Q, DC-FCCS was first performed on a negative control co-expressing Ptn-mEGFP and PMT-mApple to measure the lower limit (Q_neg_) and on a positive control expressing PMT-mEGFP-mApple to measure the upper limit (Q_pos_). Embryos coexpressing Ptn-mEGFP and Ptprz1b-mApple were measured by DC-FCCS under the same settings to calculate respective Q values.

### Phalloidin staining

Phalloidin staining was performed as described previously (Koppen *et al*, 2006). Briefly, embryos were fixed in 4% PFA/PBST at 4□ overnight. After three washes with 1X PBST for 5 min each, chorions were removed manually using forceps, and embryos were permeabilized in 2% Triton X-100/PBST at room temperature for 2 hrs. Alexa Fluor™ 488 Phalloidin methanol stock solution (A12379, ThermoFisher) was diluted 40 times in PBST according to manufacturer’s instruction and added onto embryos for staining at 4□ overnight. Stained embryos were washed three times with 1X PBST for 5 min each. After a counterstain with 5 μg/mL DAPI for 30 min, samples were mounted on imaging glass-bottom dishes with 1.5% (w/w) low melting agarose for confocal imaging.

### Fluorescent in situ hybridization

Fluorescent *in situ* hybridization (FISH) was performed using Fast Blue BB Salt hemi (zinc chloride) salt (Fast Blue) (44670, Sigma) and Naphthol AS-MX phosphate disodium salt (Naphthol) (N5000, Sigma) following a published protocol (Lauter *et al*, 2011). The Fast Blue staining signals are blue under brightfield and can be excited by 633 nm laser to emit fluorescence with a wavelength longer than 650 nm for confocal imaging on LSM900 (Zeiss).

## RT-PCR and real-time qPCR

Total RNA extractions were performed using a TRIzol-chloroform method following the manufacturer’s manual (15596026, ThermoFisher). The concentration of extracted total RNA was determined using Qubit™ RNA HS Assay Kit (Q32852, ThermoFisher) following manufacturer’s instructions. First strand cDNA synthesis was conducted using a RevertAid First Strand cDNA Synthesis Kit (K1621, ThermoFisher). Residual genomic DNA was digested by DnaseI (EN0521, ThermoFisher), and the DnaseI was deactivated by heating at 65 □ for 10 mins with 1 μL of 50 mM EDTA added to prevent RNA hydrolysis. The resulting solution served as template RNA for first strand cDNA synthesis following the manufacturer’s instruction.

Real-time quantitative PCR (qPCR) was performed using 2X PowerUp™ SYBR™ Green Master Mix (A25743, ThermoFisher) using a CFX96 Touch Real-Time PCR Detection System (Biorad) following the manufacturer’s instructions. qPCRs comprised 40 cycles and a dissociation step to assess the specificity of the primers. The Ct values were automatically calculated based on the threshold defined by the of CFX Maestro software (Biorad). For all relative qPCR measurements, the housekeeping gene β*-actin* (*actb1*) was chosen for data normalization. Three technical replicates were set up for each gene of interest (GOI), and three biological repeats were measured for each sample. The fold change was calculated by the 2 − Δ Δ^Ct^ method. The detailed equation is listed as follows:

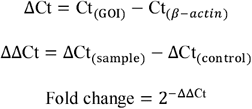

To compare the relative level of all GOI at 0 hpf, *mdka* was regarded as control for calculation. To calculate the relative fold change of *prickle* genes in MZ *mdka* mutants and MZ *ptprz1b* mutants, the expression level of WT was set as control.

### Confocal time-lapse imaging

Embryos from WT, MZ *ptprz1b*, MZ *mdka* and MZ *ptn* mutant incrosses were injected at 1-cell stage with 20 pg *PMT-mEGFP* mRNA to visualize midline or 30 pg of *H2B-EGFP* and 20 pg of *PMT-mApple* mRNA to track cell dynamics, respectively. The injected embryos were incubated until 13 hpf and mounted in 0.5% low melting agarose (Bio-rad, 1613114) in a glass bottom imaging dish. Embryos were positioned so that the dorsal hindbrain faced towards the glass bottom. Time-lapse imaging was performed on a LSM900 (Zeiss) confocal microscope. Time-lapse imaging was set up using a 63x oil lens (NA = 1.4, 0.5x zoom-out, 6 tiles-stitched) with a time interval of 4 min, 300 cycles and a Z-depth of 58 μm (2 μm, 30 slices), starting from the dorsal-most epithelium to capture midline formation. To track cell dynamics in the hindbrain, images were captured with a time interval of 3.5 mins, 120 cycles through a Z-depth of 41.17 μm (0.23 μm, 180 slices) using 63x oil lens (NA = 1.4, 0.5x zoom-out) or interval of 4 min for 160 cycles using a 20x lens (NA = 0.8) with a Z-depth of 47.17 μm (0.53 μm, 90 slices) starting from the dorsal-most epithelium.

To visualize the subcellular localization of Prickle, mRNAs encoding Drosophila Prickle-GFP (30 pg) and PMT-mApple (20 pg) were co-injected into WT, MZ *mdka*, or MZ *ptprz1b* embryos at 1-cell stage. Time-lapse imaging was performed on a LSM900 (Zeiss) confocal microscope, using a 20x lens (NA = 0.8). Images were captured with a time interval of 3.5 mins, 120 cycles through a Z-depth of 59 μm (1 μm, 60 slices).

### Cell division and tracking analysis

Cell division analysis was performed on time-lapse images from *H2B-EGFP* and *PMT-mApple* mRNA injected embryos. Maximum-intensity-projections (MIPs) were performed, and a region of interest (ROI) containing rhombomeres 1 to 4 was chosen. Periods of 70 mins for WT and MZ *ptprz1b* mutants, and 72 mins for MZ *mdka* mutants were analyzed at a time point before midline structures were visible, which roughly corresponded to 15 to 16 hpf. The directions of cell divisions were measured at the telophase stage on a XY plane. The line function of ImageJ was used to link the middle points of two separating pairs of chromosomes at telophase. Relative distances of cell divisions away from the midline were determined by measuring the distance along the mediolateral axis between the middle point position of the cell division to the presumptive midline. The midpoint position of the cell division was calculated based on the drawn line.

Cell tracking analysis was performed on cells undergoing midline-crossing C-divisions in hindbrains at stages between 15 to 17 hpf. For each embryo, cells were randomly selected at rhombomeres 1 to 4 in WT, MZ *mdka*, and MZ *ptprz1b* mutants. Nuclei centres were manually labelled using the point function in IMARIS 9.5 (Bitplane) and arbitrarily regarded as indicator of cell position. This labelling was performed on each time frame until the nuclei were no longer distinguishable or reached their predicted destination. Segmentation was conducted with the surface function of IMARIS to segregate the nucleus of interest.

### Reaction-diffusion modeling

We considered a simple reaction-diffusion model. The ligand, Ptn (denoted by *L*) is taken to be expressed at one end of the domain and diffuses through the system with diffusion constant, *D*, and degrades at rate μ. Ptn binds to its receptor, Ptprz1 (denoted by *R*), which for simplicity we assume to be expressed uniformly through the embryo. Upon binding, the receptor-ligand complex is internalised into the cell and presumed to be degraded.

If the Ptn distribution follows the basic synthesis-diffusion-decay model as given above, the steady state solution for both the wild type and control experiments can be approximated as:

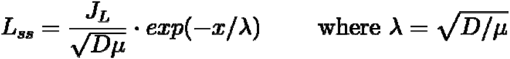

We can thus fit the experimental measurements of Ptn in the control and wildtype, with an exponential equation, *y* − *a*.*exp*(x * *b*)to obtain its decay length λ.

The average decay length of Ptn across all the experimental measurements is 570 ± 120μm (n=5, discounting one outlier). The diffusion coefficient of Ptn is estimated to be about 61 μm^2^s^-1^ in the extracellular space from FCS measurements (**Fig S1K-L**). This places the decay rate of the ligand around ∼1.6 x 10^-4^ s^-1^, or a mean lifetime of ∼1.8 hours. Since FCS gives the local diffusion coefficient and does not take into account the tortuosity of the ECS environment, it could be expected that the numbers measured represent an upper estimate of effective ptn-MYC diffusion, and the mean lifetime to be longer than given.

### Simulations

Simulations of the theoretical model of Ligand and Receptor interactions were performed on a 1D domain *x ∈*[0,400], matching the experimental distances measured in the embryo. ***L*** has a no flux boundary condition at and *x*=0and *x*=400.

*R* was initially uniformly present in the domain at the concentration

, corresponding roughly to the 1 hpf it takes to reach the 8-cell stage. The simulation was then run for 3 hours of in-simulation time, corresponding to ∼4 hpf, when immunostaining takes place. The production rate of L was held at 1 mol m^-3^ s^-1^ for all simulations. The evolution of the receptor and ligand distributions over time is plotted in **Fig S6**.

At low levels of J_R_, the receptors are consumed entirely throughout the domain, while a ligand distribution is established across the domain. In comparison, at high levels of J_R_, only the left hand side of the system is depleted of R, while the ligand distribution is confined to the left-hand side and is depleted when it enters a region of high R. This gives a very sharp transition boundary between regions with high concentrations of ligand and high concentration of receptors (**Fig S6C-C’**).

Decreasing the ligand-receptor binding rate also decreases the sharpness of the boundary between the regions of high ligand concentration and regions of high receptor concentration.

### Statistical analysis

Statistical analyses were performed using EstimationStats (https://www.estimationstats.com/#/) (Claridge-Chang & Assam, 2016; Ho *et al*, 2019). Briefly, after computational bootstrap resampling, a 95% or 99% confidence interval (CI) of the mean difference was calculated after computational bootstrap resampling. Two-sided permutation *t*-tests were performed to calculate *P*-values for each set of comparison. When *P* < 0.05 (CI = 95%) or *P* < 0.01 (CI = 99%), the set of comparison was considered to be significantly different. For statistical comparison of qPCR data, the built-in statistical function of CFX Maestro software (Biorad) was used. One-way ANOVA was conducted with a CI of 95%. To compare cell division orientation profile between WT and MZ *ptprz1b* or WT and MZ *mdka* mutants, Kolmogorov–Smirnov test was performed with a CI of 95%.

## ACKNOWLEDGEMENTS

The authors would like to thank the Department of Biological Sciences (NUS) confocal unit and fish facility staff for their constant support. The authors also thank Brian Ciruna for generously providing the *Drosophila Prickle* construct and Paul Matsudaira for the *H2B-EGFP* construct. This work was funded by grants from the Singapore Ministry of Education (grant numbers MOE-2016-T3-1-005 and MOE-T2EP30221-0008).

## AUTHOR CONTRIBUTIONS

YL, TES, TW and CW conceived and designed the research. YL, KR and TYJL performed the experiments under supervision of TES, TW and C.W.

## REFERENCES

Amet LE, Lauri SE, Hienola A, Croll SD, Lu Y, Levorse JM, Prabhakaran B, Taira T, Rauvala H, Vogt TF (2001) Enhanced hippocampal long-term potentiation in mice lacking heparin-binding growth-associated molecule. Mol Cell Neurosci 17: 1014–1024

Bai Y, Tan X, Zhang H, Liu C, Zhao B, Li Y, Lu L, Liu Y, Zhou J (2014) Ror2 receptor mediates Wnt11 ligand signaling and affects convergence and extension movements in zebrafish. J Biol Chem 289: 20664–20676

Bhaduri A, Di Lullo E, Jung D, Muller S, Crouch EE, Espinosa CS, Ozawa T, Alvarado B, Spatazza J, Cadwell CR et al (2020) Outer Radial Glia-like Cancer Stem Cells Contribute to Heterogeneity of Glioblastoma. Cell Stem Cell 26: 48–63 e46

Buckley CE, Ren X, Ward LC, Girdler GC, Araya C, Green MJ, Clark BS, Link BA, Clarke JD (2013) Mirror-symmetric microtubule assembly and cell interactions drive lumen formation in the zebrafish neural rod. EMBO J 32: 30–44

Calinescu AA, Vihtelic TS, Hyde DR, Hitchcock PF (2009) Cellular expression of midkine-a and midkine-b during retinal development and photoreceptor regeneration in zebrafish. J Comp Neurol 514: 1–10

Chang MH, Huang CJ, Hwang SP, Lu IC, Lin CM, Kuo TF, Chou CM (2004) Zebrafish heparin-binding neurotrophic factor enhances neurite outgrowth during its development. Biochem Biophys Res Commun 321: 502–509

Ciruna B, Jenny A, Lee D, Mlodzik M, Schier AF (2006) Planar cell polarity signalling couples cell division and morphogenesis during neurulation. Nature 439: 220–224

Claridge-Chang A, Assam PN (2016) Estimation statistics should replace significance testing. Nature Methods 13: 108–109

Clarke J (2009) Role of polarized cell divisions in zebrafish neural tube formation. Curr Opin Neurobiol 19: 134–138

Concha ML, Adams RJ (1998) Oriented cell divisions and cellular morphogenesis in the zebrafish gastrula and neurula: a time-lapse analysis. Development 125: 983–994

Fan QW, Muramatsu T, Kadomatsu K (2000) Distinct expression of midkine and pleiotrophin in the spinal cord and placental tissues during early mouse development. Dev Growth Differ 42: 113–119

Foo YH, Korzh V, Wohland T (2011) Fluorescence Correlation and Cross-Correlation Spectroscopy Using Fluorescent Proteins for Measurements of Biomolecular Processes in Living Organisms. In: Fluorescent Proteins II, pp. 213–248.

Harroch S, Palmeri M, Rosenbluth J, Custer A, Okigaki M, Shrager P, Blum M, Buxbaum J, Schlessinger J (2000) No obvious abnormality in mice deficient in receptor protein tyrosine phosphatase β. Molecular and cellular biology 20: 7706–7715

Hirano S, Mii Y, Charras G, Michiue T (2022) Alignment of the cell long axis by unidirectional tension acts cooperatively with Wnt signalling to establish planar cell polarity. Development 149

Ho J, Tumkaya T, Aryal S, Choi H, Claridge-Chang A (2019) Moving beyond P values: data analysis with estimation graphics. Nature Methods 16: 565–566

Hong E, Brewster R (2006) N-cadherin is required for the polarized cell behaviors that drive neurulation in the zebrafish. Development 133: 3895–3905

Kimmel CB, Ballard WW, Kimmel SR, Ullmann B, Schilling TF (1995) Stages of embryonic development of the zebrafish. Developmental dynamics 203: 253–310

Koppen M, Fernandez BG, Carvalho L, Jacinto A, Heisenberg CP (2006) Coordinated cell-shape changes control epithelial movement in zebrafish and Drosophila. Development 133: 2671–2681

Kruse K, Pantazis P, Bollenbach T, Julicher F, Gonzalez-Gaitan M (2004) Dpp gradient formation by dynamin-dependent endocytosis: receptor trafficking and the diffusion model. Development 131: 4843–4856

Kuboyama K, Fujikawa A, Suzuki R, Noda M (2015) Inactivation of Protein Tyrosine Phosphatase Receptor Type Z by Pleiotrophin Promotes Remyelination through Activation of Differentiation of Oligodendrocyte Precursor Cells. J Neurosci 35: 12162–12171

Kuhn T, Landge AN, Morsdorf D, Cossmann J, Gerstenecker J, Capek D, Muller P, Gebhardt JCM (2022) Single-molecule tracking of Nodal and Lefty in live zebrafish embryos supports hindered diffusion model. Nat Commun 13: 6101

Lauter G, Söll I, Hauptmann G (2011) Two-color fluorescent in situ hybridization in the embryonic zebrafish brain using differential detection systems. BMC Developmental Biology 11: 43

Ma X, Foo YH, Wohland T (2014) Fluorescence cross-correlation spectroscopy (FCCS) in living cells. Methods Mol Biol 1076: 557–573

Maeda N, Ichihara-Tanaka K, Kimura T, Kadomatsu K, Muramatsu T, Noda M (1999) A Receptor-like Protein-tyrosine Phosphatase PTPζ/RPTPβ Binds a Heparin-binding Growth Factor Midkine INVOLVEMENT OF ARGININE 78 OF MIDKINE IN THE HIGH AFFINITY BINDING TO PTPζ. Journal of Biological Chemistry 274: 12474–12479

Maeda N, Nishiwaki T, Shintani T, Hamanaka H, Noda M (1996) 6B4 proteoglycan/phosphacan, an extracellular variant of receptor-like protein-tyrosine phosphatase ζ/RPTPβ, binds pleiotrophin/heparin-binding growth-associated molecule (HB-GAM). Journal of Biological Chemistry 271: 21446–21452

Marlow F, Topczewski J, Sepich D, Solnica-Krezel L (2002) Zebrafish Rho kinase 2 acts downstream of Wnt11 to mediate cell polarity and effective convergence and extension movements. Curr Biol 12: 876–884

Matsubara S, Take M, Pedraza C, Muramatsu T (1994) Mapping and characterization of a retinoic acid-responsive enhancer of midkine, a novel heparin-binding growth/differentiation factor with neurotrophic activity. J Biochem 115: 1088–1096

Meng K, Rodriguez-Pena A, Dimitrov T, Chen W, Yamin M, Noda M, Deuel TF (2000) Pleiotrophin signals increased tyrosine phosphorylation of beta -catenin through inactivation of the intrinsic catalytic activity of the receptor-type protein tyrosine phosphatase beta /zeta. Proceedings of the National Academy of Sciences 97: 2603–2608

Muramatsu H, Zou P, Kurosawa N, Ichihara-Tanaka K, Maruyama K, Inoh K, Sakai T, Chen L, Sato M, Muramatsu T (2006) Female infertility in mice deficient in midkine and pleiotrophin, which form a distinct family of growth factors. Genes Cells 11: 1405–1417

Nakamura E, Kadomatsu K, Yuasa S, Muramatsu H, Mamiya T, Nabeshima T, Fan Q-W, Ishiguro K, Igakura T, Matsubara S et al (1998) Disruption of the midkine gene (Mdk) resulted in altered expression of a calcium binding protein in the hippocampus of infant mice and their abnormal behaviour. 3: 811–812

Olmeda D, Cerezo-Wallis D, Riveiro-Falkenbach E, Pennacchi PC, Contreras-Alcalde M, Ibarz N, Cifdaloz M, Catena X, Calvo TG, Canon E et al (2017) Whole-body imaging of lymphovascular niches identifies pre-metastatic roles of midkine. Nature 546: 676–680

Pollen AA, Nowakowski TJ, Chen J, Retallack H, Sandoval-Espinosa C, Nicholas CR, Shuga J, Liu SJ, Oldham MC, Diaz A et al (2015) Molecular identity of human outer radial glia during cortical development. Cell 163: 55–67

Qin EY, Cooper DD, Abbott KL, Lennon J, Nagaraja S, Mackay A, Jones C, Vogel H, Jackson PK, Monje M (2017) Neural Precursor-Derived Pleiotrophin Mediates Subventricular Zone Invasion by Glioma. Cell 170: 845–859 e819

Quesada-Hernandez E, Caneparo L, Schneider S, Winkler S, Liebling M, Fraser SE, Heisenberg CP (2010) Stereotypical cell division orientation controls neural rod midline formation in zebrafish. Curr Biol 20: 1966–1972

Rohrschneider MR, Elsen GE, Prince VE (2007) Zebrafish Hoxb1a regulates multiple downstream genes including prickle1b. Dev Biol 309: 358–372

Schafer M, Rembold M, Wittbrodt J, Schartl M, Winkler C (2005) Medial floor plate formation in zebrafish consists of two phases and requires trunk-derived Midkine-a. Genes Dev 19: 897–902

Schneider CA, Rasband WS, Eliceiri KW (2012) NIH Image to ImageJ: 25 years of image analysis. Nature Methods 9: 671–675

Shi Y, Ping YF, Zhou W, He ZC, Chen C, Bian BS, Zhang L, Chen L, Lan X, Zhang XC et al (2017) Tumour-associated macrophages secrete pleiotrophin to promote PTPRZ1 signalling in glioblastoma stem cells for tumour growth. Nat Commun 8: 15080

Shintani T, Watanabe E, Maeda N, Noda M (1998) Neurons as well as astrocytes express proteoglycan-type protein tyrosine phosphatase zeta/RPTPbeta: analysis of mice in which the PTPzeta/RPTPbeta gene was replaced with the LacZ gene. Neurosci Lett 247: 135–138

Soderberg O, Gullberg M, Jarvius M, Ridderstrale K, Leuchowius KJ, Jarvius J, Wester K, Hydbring P, Bahram F, Larsson LG et al (2006) Direct observation of individual endogenous protein complexes in situ by proximity ligation. Nat Methods 3: 995–1000

Tawk M, Araya C, Lyons DA, Reugels AM, Girdler GC, Bayley PR, Hyde DR, Tada M, Clarke JD (2007) A mirror-symmetric cell division that orchestrates neuroepithelial morphogenesis. Nature 446: 797–800

Tomomura M, Kadomatsu K, Matsubara S, Muramatsu T (1990) A retinoic acid-responsive gene, MK, found in the teratocarcinoma system. Heterogeneity of the transcript and the nature of the translation product. Journal of Biological Chemistry 265: 10765–10770

van Eekelen M, Overvoorde J, van Rooijen C, den Hertog J (2010) Identification and expression of the family of classical protein-tyrosine phosphatases in zebrafish. PLoS One 5: e12573

Wang Y, Wang X, Wohland T, Sampath K (2016) Extracellular interactions and ligand degradation shape the nodal morphogen gradient. Elife 5

Weicksel SE, Gupta A, Zannino DA, Wolfe SA, Sagerstrom CG (2014) Targeted germ line disruptions reveal general and species-specific roles for paralog group 1 hox genes in zebrafish. BMC Dev Biol 14: 25

Winkler C, Moon R (2001a) Zebrafish mdk2, a novel secreted midkine, participates in posterior neurogenesis. Developmental Biology 229: 102–118

Winkler C, Moon RT (2001b) Zebrafish mdk2, a novel secreted midkine, participates in posterior neurogenesis. Dev Biol 229: 102–118

Winkler C, Schafer M, Duschl J, Schartl M, Volff JN (2003) Functional divergence of two zebrafish midkine growth factors following fish-specific gene duplication. Genome Res 13: 1067–1081

Winkler C, Yao S (2014) The midkine family of growth factors: diverse roles in nervous system formation and maintenance. Br J Pharmacol 171: 905–912

Yao S, Cheng M, Zhang Q, Wasik M, Kelsh R, Winkler C (2013) Anaplastic lymphoma kinase is required for neurogenesis in the developing central nervous system of zebrafish. PLoS One 8: e63757

Zhu S, Liu L, Korzh V, Gong Z, Low BC (2006) RhoA acts downstream of Wnt5 and Wnt11 to regulate convergence and extension movements by involving effectors Rho kinase and Diaphanous: use of zebrafish as an in vivo model for GTPase signaling. Cell Signal 18: 359–372

