## Supplementary information text for "Midkine and Ptprz1b act upstream of Wnt planar cell polarity to establish a midline in the developing zebrafish hindbrain"

### FIGURE LEGENDS

#### Figure S1. *In vivo* proximity ligation assay of zebrafish Mdka, Mdkb, Ptn and Ptprz1a and representatiFve FCCS and FCS measurements.

(**A**) Principle of proximity ligation assay (PLA). When Mdk/Ptn-MYC and HA-Ptprz1 interact, close physical distance allows pairing of (+) and (-) oligonucleotides on secondary antibodies to generate DNA concatemeric barcodes for Texas Red (TR)-labelled probes to bind. Ptprz1 consists of a carbonic anhydrase (CA), a fribronectin-3 (FN3) and two intracellular protein tyrosine phosphatase c domains (PTPc). Chondroitin sulfate (CS) side chains are illustrated as dashed lines.

(**B**) Experimental setup of *in vivo* PLA in zebrafish embryos. mRNAs encoding Mdk/Ptn-MYC and HA-Ptprz1 were co-injected in WT embryos at 1-cell stage. Injected embryos were fixed at 80% epiboly, subjected to PLA and subsequent confocal imaging.

(**C**) Representative maximum-intensity-projected (MIP) PLA images of Mdka-Ptprz1a, Mdkb-Ptprz1a and Ptn-Ptprz1a pairs. PLA signals are coloured in cyan, DAPI in magenta. Scale bars = 20 μm.

(**D-F**) Statistical analysis of PLA levels normalized to DAPI signals after thresholding (PLA/DAPI area ratios). Data are shown as dots with mean ± SD indicated. Data of Mdka+S-EGFP, Mdka+Ptprz1b, Mdkb+Ptprz1b and Ptn-Ptprz1b are also shown in **Fig.1E**. Statistical analysis was conducted using Estimation Stats (<https://www.estimationstats.com>) to compare the mean of each dataset to that of a random collision control (**E**), the mean of Mdk/Ptn-Ptprz1a and that of Mdk/Ptn-Ptprz1b, the mean of Mdka-Ptprz1b, Mdkb-Ptprz1b and Ptn-Ptprz1b (**F**), or the mean of Mdka-Ptprz1a, Mdkb-Ptprz1a and Ptn-Ptprz1a (**F**). *P* values are determined using a two-sided permutation *t*-test with a CI of 95%. Statistical significance (*P* < 0.05) is indicated by asterisks.

(**G-J**) FCCS measurements were performed on MZ *ptprz1b* embryos at 4 hpf that had been injected with the indicated mRNAs at 1-cell stage. Green and red curves are auto-correlation functions plotted from green and red fluorophore channels, respectively. Blue curves represent the cross-correlation function. All curves were fitted by a 3D-2 particles model as represented by black curves, respectively. **(G)** MZ *ptprz1b* mutant embryos expressing Ptn-mEGFP and PMT-mApple were used as negative control. **(H)** MZ *ptprz1b* mutant embryos expressing membrane bound PMT-mEGFP-mApple were used as positive control. **(I)** Data obtained from MZ *ptprz1b* embryos that were co-injected with *ptn-mEGFP* and *ptprz1b-mApple* mRNA. The relative cross-correlation values of FCCS measurements are shown in (**J**) in the form of mean ± SD.

(**K**) Representative FCS measurement on WT embryos at 4hpf injected with *ptn-mEGFP* at 1-cell stage. Green curve are auto-correlation functions, which were fitted by a 3D-2 particles model as shown by black curve. The diffusion coefficient of fast components is shown in (**L**) as mean ± SD.

#### Figure S2. Generation of *mdka*, *mdkb*, *ptn* and *ptprz1b* mutants.

(**A**) Gene locus organization for *mdka*, *mdkb*, *ptn* and *ptprz1b*. Black rectangular boxes represent 5’ or 3’ UTRs, while boxes in magenta indicate CDS. gRNA targeting sites are indicated by perpendicular arrows. Primer binding sites for genotyping are shown as horizontal arrows. For *ptprz1b*, CRISPR target sites were selected to delete the entire transcribed region and avoid genetic compensation (Rossi, et al., 2015). The generated *mdka* and *ptn* alleles produced stable mRNAs that were predicted to encode truncated proteins by RT-PCR with Phusion^TM^ DNA Polymerase (F530S, ThermoFisher), electrophoresis and sequencing with BigDyeTM Terminator 3.1 (4337455, ThermoFisher) where applicable.

(**B**) RT-PCR of mutant *mdka*, *mdkb*, and *ptn* transcripts in maternal-zygotic (MZ) mutants. The full length CDS region was amplified in WT and corresponding mutants at 24 hpf, respectively. (**C-D**) Mutant Mdka and Ptn protein sequences were predicted based on sequenced mutant *mdka* and *ptn* mRNA transcripts.

(**C**) NCBI blast alignment of WT and mutant Mdka amino acid sequences. Signal peptide, N-domain and C-domain sequences are indicated by blue, magenta and cyan boxes, respectively.

(**D**) NCBI blast alignment of WT and mutant Ptn amino acid sequences. Blue, magenta and cyan boxes label signal peptide, N-domain and C-domain sequences of Ptn, respectively.

#### Figure S3. Severest rhombomeric midline phenotype in MZ *ptprz1b*, MZ *mdka* and MZ *ptn* mutants and disorganized nuclei distribution in MZ *ptprz1b* and MZ *mdka* mutants.

(**A-E**) Confocal time-lapse images showing severe midline defects in MZ *ptprz1b* (**A-B**), MZ *mdka* (**C**) and MZ *ptn* mutants (**D**). Embryos were imaged from approx. 13 hpf with 4 min intervals with PMT-mEGFP (**A-B, D**) or PMT-mApple (**C**) to label plasma membranes. (**A-D**) Representative MIP images (dorsal views) at 4 and 6 h of time-lapse. Elapsed time (*t*) is shown in the form of hours:minutes. (**E**) Reconstructed orthogonal view of r2 in MZ *ptprz1b* mutant. Yellow lines label midline structures in (**A’-E’**). Solid magenta line in (**A’**) indicates position of views shown in (**E, E’**). Dotted magenta lines in (**B’**) delineate rhombomere boundaries. Asterisks label position of otic vesicles, adjacent to r5. Scale bars = 50 μm.

(**F-H**) Representative MIP images of Phalloidin and DAPI-stained MZ *ptprz1b*, MZ *mdka* and MZ *ptn* mutant embryos with severest phenotype at 17 hpf. Phalloidin-stained F-actin is pseudo-coloured in cyan, and DAPI-stained nuclei in magenta. Asterisks label positions of otic vesicles. Yellow boxes indicate regions shown with higher magnification in (**F’-H’**). F-actin aggregation is highlighted with arrowheads in (**F’-H’**). Scale bars = 50 μm (**F-H**) and 30 μm (**F’-H’**).

(**I-K**) Mean fluorescent intensity histograms of Phalloidin (cyan) and DAPI (magenta) along mediolateral axis of (**F’-H’**). Asterisks show accumulation of Phalloidin-stained F-actin.

(**L-N**) Reconstructed orthogonal views of DAPI-stained nuclei (in magenta). Yellow lines delineate outer surface and midline of neural keel. White lines indicate paths used for measurement of fluorescent intensity as shown in (**O-Q**). Scale bars = 20 μm.

(**O-Q**) Respective mean fluorescent intensity histogram of DAPI (magenta) along axis indicated in (**L-N**).

**Figure S4. Delayed midline formation and accelerated neural keel convergence in MZ *mdkb* mutants.**

(**A-B**) Representative MIP images of Phalloidin (cyan) and DAPI (magenta) stained WT and MZ *mdkb* mutant embryos at 17 hpf. WT showed distinctive F-actin accumulation in medial region (**A**), in contrast to MZ *mdkb* mutants (**B**). Nuclei distribution was similar in WT and MZ *mdkb* mutants (lower images). Positions of otic vesicles are labelled by white asterisks. Yellow boxes indicate regions shown with higher magnification in (**A’-B’**). Scale bars = 50 μm (**A-B**) and 30 μm (**A’-B’**).

(**C-D**) Quantification of mean fluorescent intensity of Phalloidin (cyan) and DAPI (magenta) along mediolateral axis of (**A’-B’**). Asterisks show the accumulation of Phalloidin-stained F-actin.

(**E-F**) At 18 hpf, distinctive F-actin accumulation was found in both WT and MZ *mdkb* mutant embryos. Positions of otic vesicles are indicated by asterisks. Scale bars = 50 μm.

(**G**) Representative confocal single-plane images (dorsal views) of Phalloidin-stained hindbrains in 15 hpf WT (N = 8) and MZ *mdkb* (N = 8) embryos. Asterisks indicate the positions of otic vesicles, and dotted lines mark positions of orthogonal views shown in (**H**). Scale bars = 50 μm.

(**H**) Reconstructed orthogonal views of embryos shown in (**G**). Phalloidin-stained F-actin is pseudo-coloured in cyan, and DAPI-stained nuclei in magenta. Scale bars = 50 μm.

(**I**) Quantification of the maximum width measured for individual r2, r3 and r4. Data are shown as scattered dots. Mean ± SD of each dataset is shown by line and error bar. Statistical comparison was performed between WT (N = 8) and MZ *mdkb* (N = 8) on Estimation Stats (https://www.estimationstats.com). *P* values was calculated by a two-sided permutation *t*-test performed under a CI of 95%. Statistical significance (*P* < 0.05) is indicated by asterisk.

#### Figure S5. Diffusion of Ptn affects distribution of membrane-localized Ptprz1b.

(**A**) Representative single channel MIP images from diffusion assays of Ptn-MYC. The experimental setup is described in (**Fig 3J)**. Ptn-MYC (magenta) was secreted from cells on the left and diffused to the right. Fluorescent signals from mEGFP-Ptprz1b (green), encoded by mRNA injected at 1-cell stage, is reduced on left side. Scale bar = 20 μm.

(**B**) Distribution of PMT-mEGFP control. Scale bar = 20 μm.

#### Figure S6. Reaction-diffusion model is consistent with the *in vivo* measurements.

(**A-A’**) Measured Ptn-Myc profile with fit to exponential gradient to extract decay length (solid lines) under different conditions. Each color represents a different embryo.

(**B-B’**) Corresponding model simulations (Methods) D = 30 µm^2^ s^-1^, µ = 2.8 × 10^-4^ s^-1^, J_L_ = 1 mol µm ^-1^ s^-1^).

(**C-C’**) Parameter exploration in α-J_R_ space (D = 30 µm^2^ s^-1^, µ = 2.8 × 10^-5^ s^-1^, J_L_ = 1 mol µm ^-1^ s^-1^). There exists a switching in behavior of the ligand-receptor concentration at intermediate parameter values. See Methods for details of equations and simulation environment.

#### Figure S7. Maternal loss of *ptprz1b* but not *mdka* causes midline defects in rhombomeres.

(**A-D**) Representative MIP images of Phalloidin (cyan) and DAPI (magenta) stained rhombomeres in 17 hpf embryos from WT incross (**A**), MZ *ptprz1b* female crossed with WT male (**B**; M *ptprz1b*), MZ *ptprz1b* male crossed with WT female (**C**), and MZ *mdka* female crossed with WT male (**D**; M *mdka*). Arrowheads label the midline. Asterisks indicate position of otic vesicles. Scale bars = 50 μm.

(**E**) Real-time qPCR showing high maternal mRNA contribution of *ptprz1b* but not of *mdka*. The relative mRNA levels of *ptprz1b* are the fold changes normalized to the level of *mdka* at 0 hpf. Error bars represent SD for each dataset. One-way ANOVA was conducted with a CI of 95% by built-in algorithm of CFX Maestro. Asterisk indicates *P* < 0.05 under a CI of 95%.

(**F**) Fluorescent *in situ* hybridization (FISH) of *ptprz1b* in 14 hpf WT embryo. Representative MIP image is presented as dorsal view and pseudo-coloured in fire LUT. Dotted lines delineate the lateral edges of rhombomeres, and asterisks indicate position of otic vesicles. Scale bar = 50 μm.

(**G**) Mean fluorescent intensity of FISH signals for *ptprz1b* as shown in (**F**) and *mdka* shown in **Fig 3B** along the anteroposterior axis of rhombomeres.

### MOVIE CAPTIONS

#### Movie S1. Midline establishment in WT rhombomeres.

The development of hindbrain rhombomeres in WT. Confocal time-lapse imaging started at 13 hpf with 4 min intervals. The plasma membrane is visualized by PMT-mEGFP labelling. A single midline emerged after 4 h of imaging.

#### Movie S2. Mild ectopic midline phenotype in MZ *ptprz1b* rhombomeres.

Representative MZ *ptprz1b* mutant that developed two transient ectopic midlines in r2 to r7. Time-lapse imaging started at 13 hpf with 4 min intervals. The plasma membrane is labelled by PMT-mEGFP. Ectopic midlines emerged after 3.5 h of imaging and gradually merged medially into one.

#### Movie S3. Severe ectopic midline phenotype in MZ *ptprz1b* rhombomeres.

Representative MZ *ptprz1b* mutant that showed two permanent ectopic midlines in r1 to r7. Time-lapse imaging started at 13 hpf with 4 min intervals. The plasma membrane is indicated by PMT-mEGFP. Embryo developed ectopic midline structures after 3 h of imaging, and midlines failed to merge at the end of imaging.

#### Movie S4. Mild ectopic midline phenotype in MZ *mdka* rhombomeres.

Representative MZ *mdka* mutant that transiently established two ectopic midlines in r2 to r6. Time-lapse imaging started at 13 hpf with 4 min intervals. The plasma membrane is labelled with PMT-mApple. After 4 h of imaging, two ectopic midlines emerged and gradually merged into one.

#### Movie S5. Severe ectopic midline phenotype in MZ *mdka* rhombomeres.

Representative MZ *mdka* mutant that formed two permanent ectopic midlines in r2 and r3. Time-lapse imaging started at 13 hpf with 4 min intervals. The plasma membrane is labelled with PMT-mApple. After 4 h of imaging, two ectopic midlines emerged in r2 to r5. At r4 and r5, midlines gradually merged into one. At r2 and r3, the midlines failed to merge and gave rise to two ventricles with an ectopic cell mass in the middle.

#### Movie S6. Representative C-division in a WT neural keel.

Trajectories and segmented nuclei of a C-division event in WT. The parent cell is coloured in magenta, and the daughter cells in cyan and green, respectively. The background shows the membrane-bound PMT-mApple signals coloured in gray and delineates the morphology of the midline. Note that the green daughter cell crosses the midline.

#### Movie S7. Representative C-division in a MZ *ptprz1b* mutant neural keel.

Trajectories and segmented nuclei of a C-division event in a MZ *ptprz1b* mutant embryo. The parent cell is coloured in magenta, and the daughter cell in cyan. After C-division, the daughter cell spent an extended period in the middle of the neural keel as shown by the cyan trajectory. Eventually, after a delay, the cell intercalated into the contralateral half of the neural keel (green trajectory). This trajectory pattern coincides with the merging of the two ectopic midlines. Background shows signals from PMT-mApple, which indicates the midline morphology.

#### Movie S8. Representative C-division in a MZ *mdka* mutant neural keel.

Trajectories and segmented nuclei of a C-division in the neural keel of a MZ *mdka* mutant embryo. The parent cell is coloured in magenta, and the daughter cell in cyan. After C-division, the daughter cell was unable to intercalate into the contralateral half of the neural keel as shown by the cyan trajectory. This cell eventually intercalated into the opposite half of the neural keel as shown by the green trajectory, when the two ectopic midlines were merging. PMT-mApple coloured in gray serves as an indicator of midline morphology.
