## Supplementary figures and images for "Midkine and Ptprz1b act upstream of Wnt planar cell polarity to establish a midline in the developing zebrafish hindbrain"

### Supplemental Figure S1

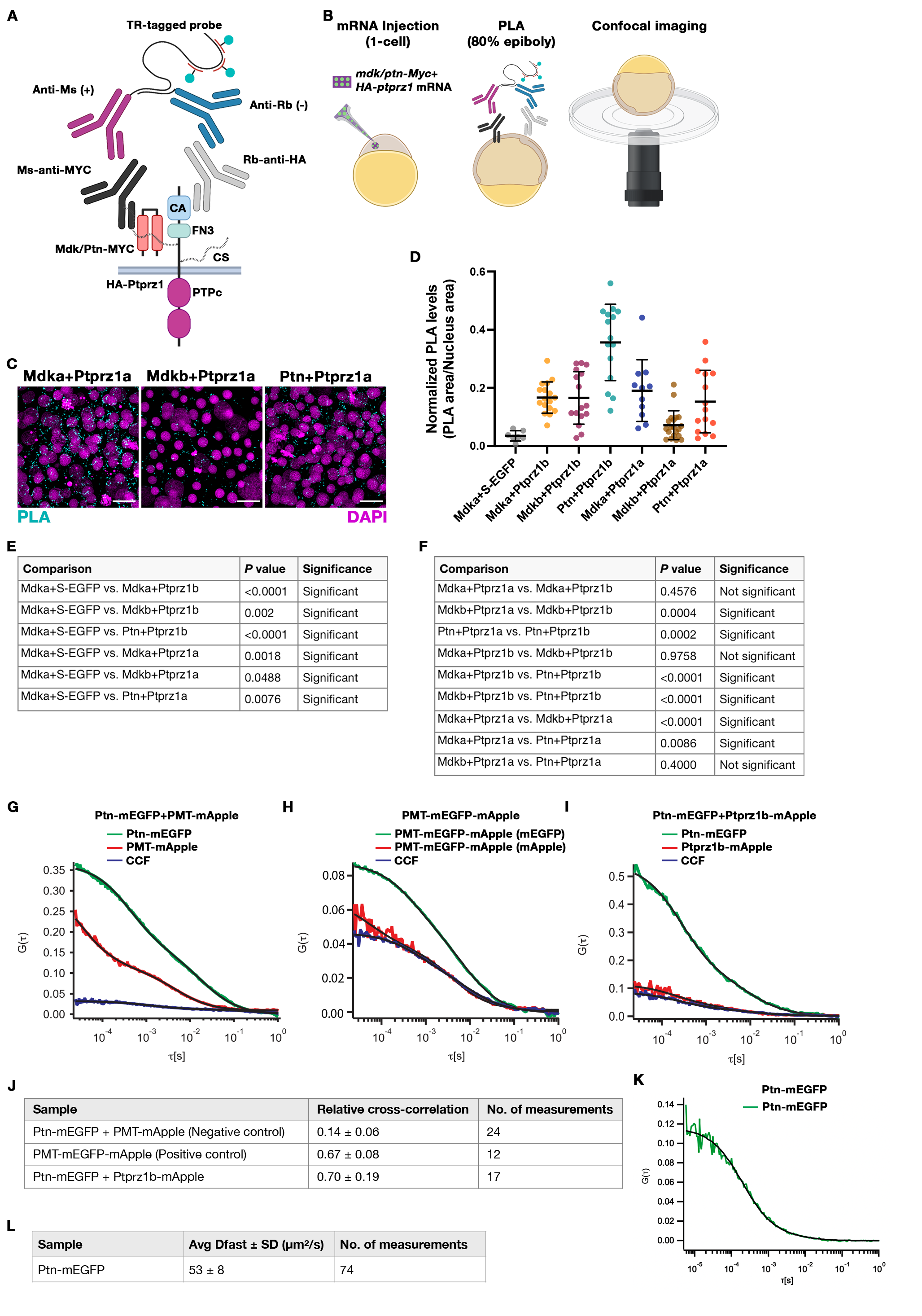

### Supplemental Figure S2

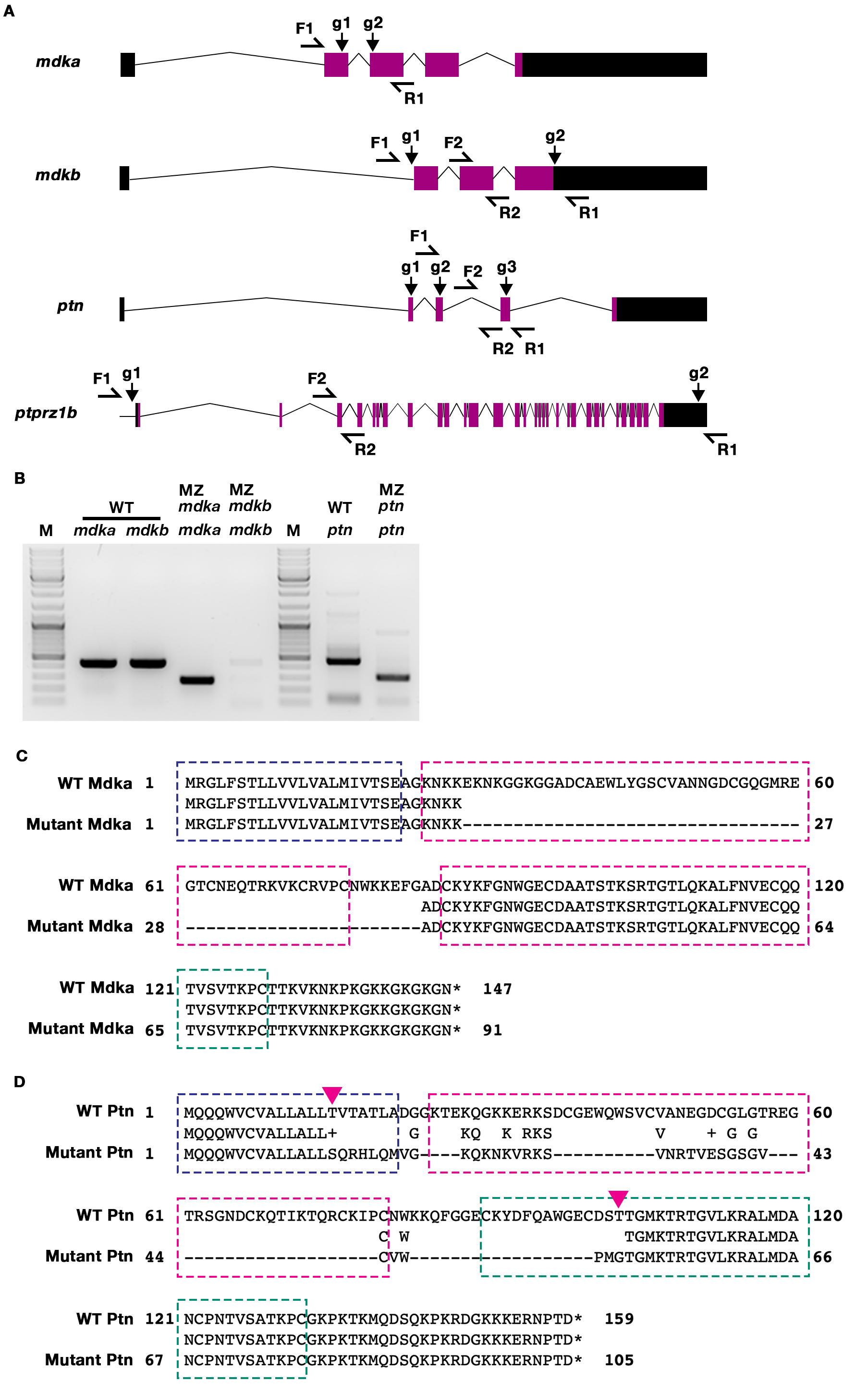

### Supplemental Figure S3

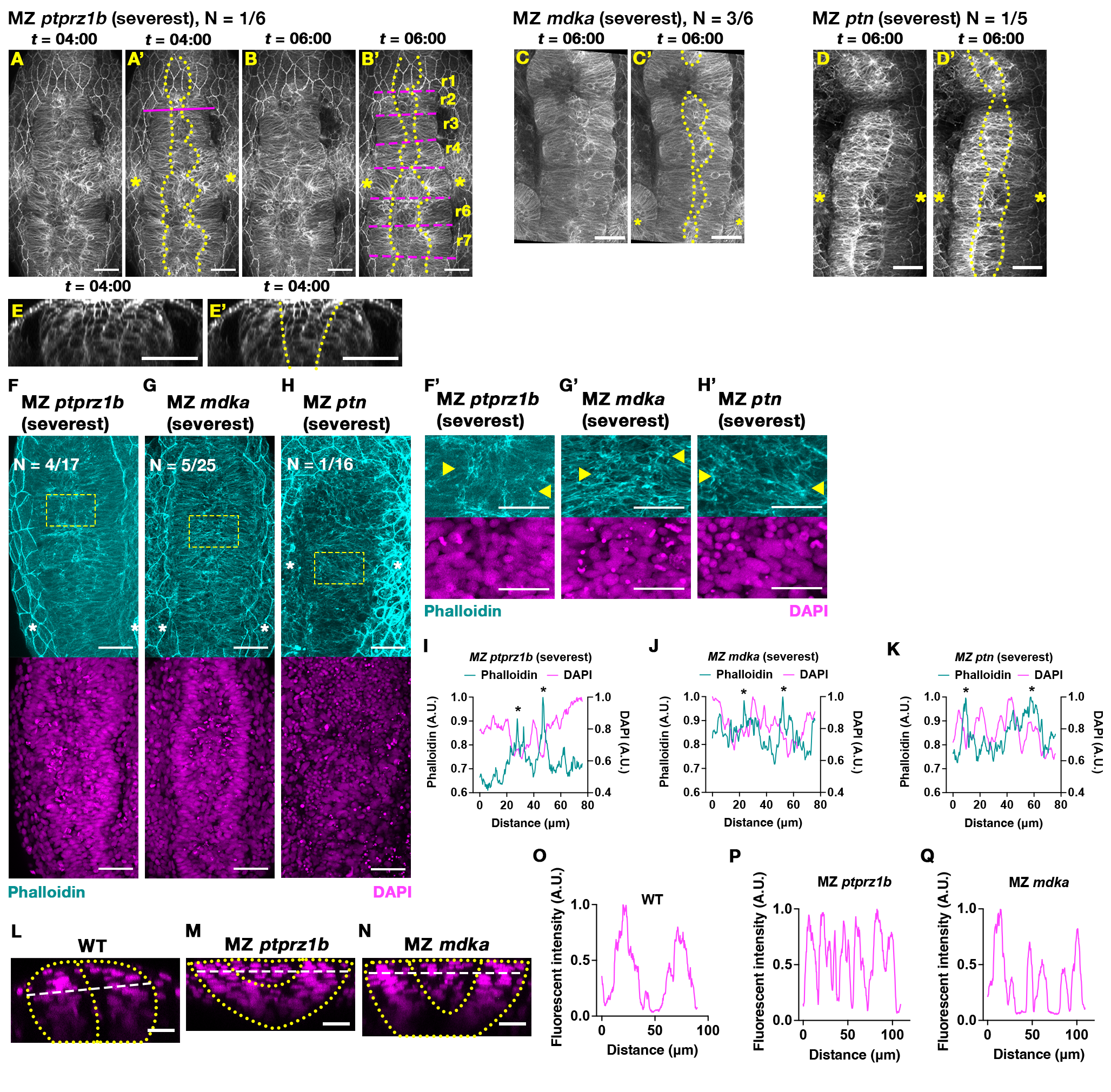

### Supplemental Figure S4

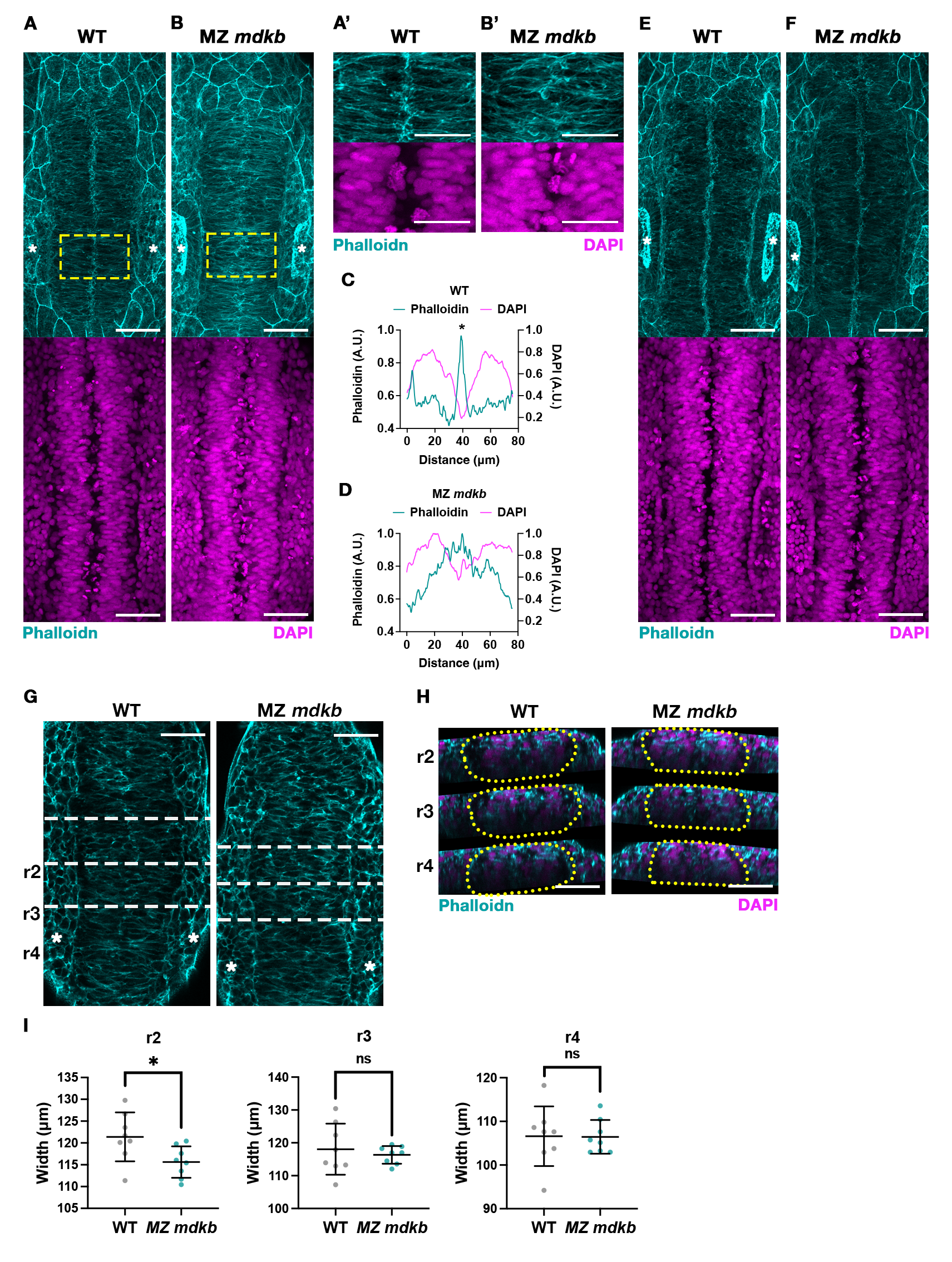

### Supplemental Figure S5

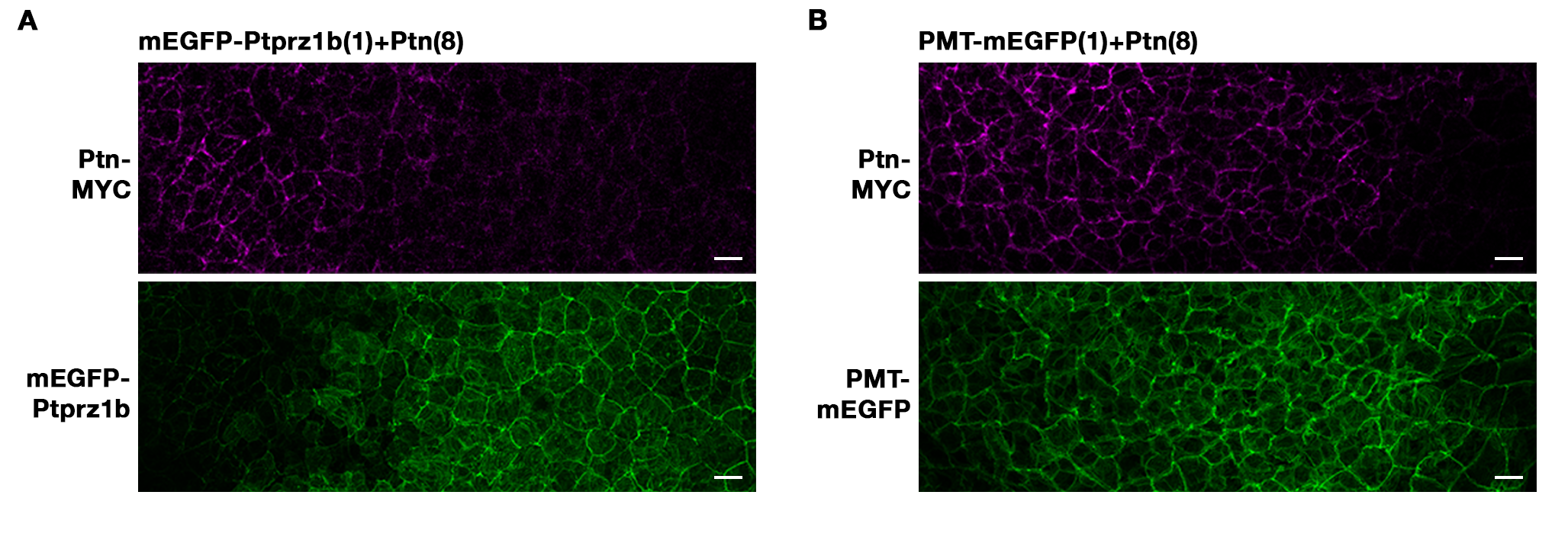

### Supplemental Figure S6

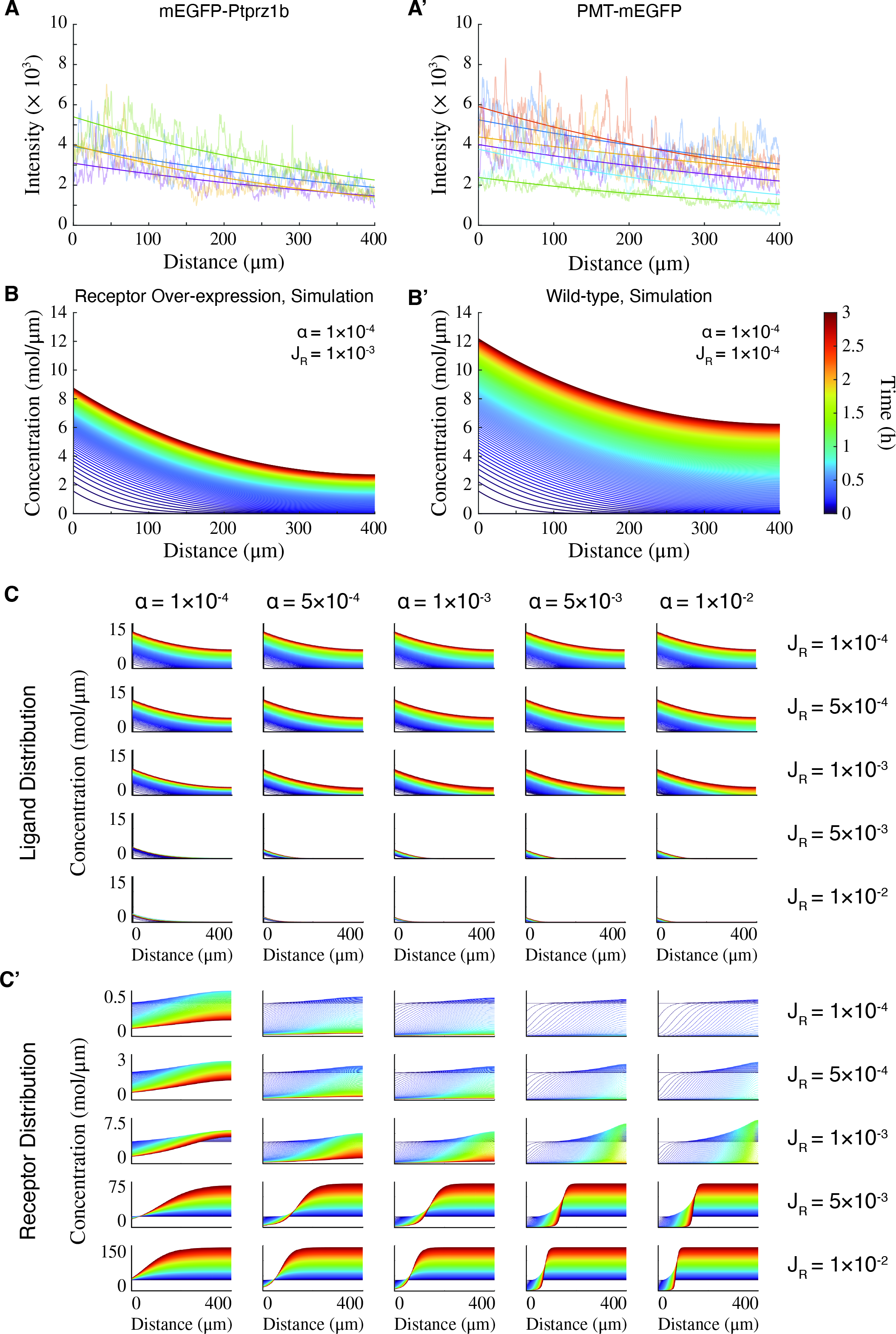

### Supplemental Figure S7

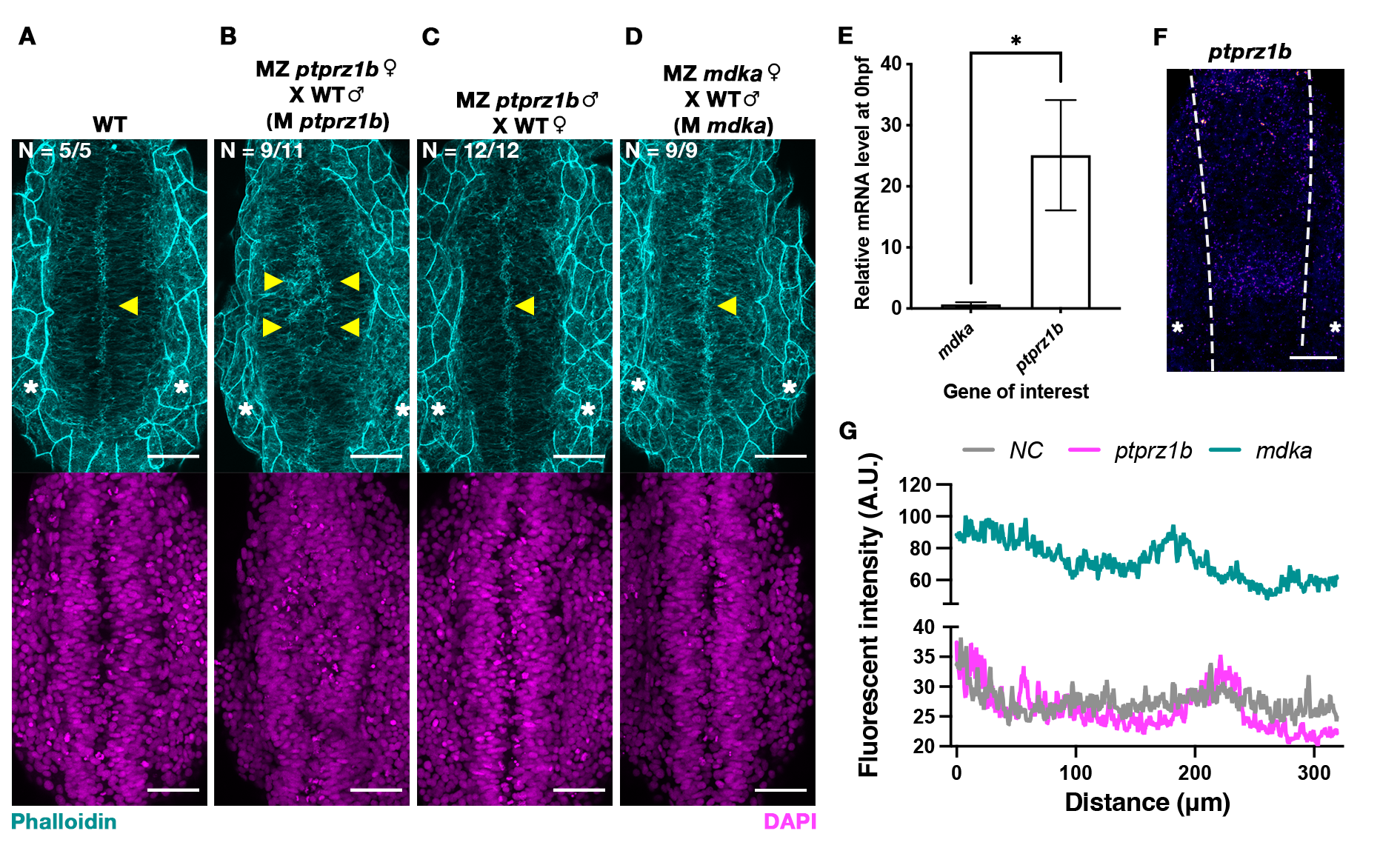
